# OPUS-DSD2: Disentangling Dynamics and Compositional Heterogeneity for Cryo-EM/ET

**DOI:** 10.1101/2024.09.24.614854

**Authors:** Zhenwei Luo, Xiangru Chen, Yiqiu Zhang, Qinghua Wang, Jianpeng Ma

## Abstract

Cryo-electron microscopy and tomography (cryo-EM/ET) capture structural heterogeneities in macromolecules, ranging from dynamic motions to compositional changes— key to understanding biological mechanisms. While the deep learning framework, OPUS-DSD, advanced heterogeneity analysis for cryo-EM, it conflates different types of heterogeneities, and remains incompatible with cryo-ET which can elucidate macromolecular functions in their native cellular environments. Here, we present OPUS-DSD2, a unified framework for disentangling structural heterogeneity in both cryo-EM and cryo-ET data. OPUS-DSD2 augments a 3D convolutional neural network with a multi-layer perceptron based rigid-body dynamics model, advocating the separation of subunit level rigid-body dynamics from other heterogeneities. Tests on real datasets demonstrate that OPUS-DSD2 effectively captures large-scale subunit motions while isolating spatially localized structural variations into distinct principal components of the composition latent space. Critically, OPUS-DSD2 enables direct analysis of noisy cryo-ET template-matching results, bypassing labor-intensive subtomogram classification and averaging and unlocking high-throughput visual proteomics. Backward-compatible with OPUS-DSD, OPUS-DSD2 is available at https://github.com/alncat/opusDSD.

## Introduction

Cryo-electron microscopy (cryo-EM) and tomography (cryo-ET) have revolutionized structural biology by enabling high resolution imaging of macromolecular complexes without crystallization^1,2^. Cryo-EM excels in high-throughput determination of purified complexes^1,3^, while cryo-ET uniquely visualizes macromolecules in their native cellular environments^4,5^, capturing spatial interactions (“molecular sociology”)^6^ and transient functional states^2^ which are critical to understanding biological mechanisms.

Traditional algorithms for structural determination in cryo-EM/ET focus on reconstructing a consensus 3D model by aggregating structural information from multiple particles in the sample. These alignment algorithms, i.e., consensus refinement in cryo-EM analysis^7^, and template matching^8^ and subtomogram averaging^4,8^ in cryo-ET analysis, determine the pose parameters of particles relative to a common model. However, the inherent structural heterogeneity in macromolecular complexes presents significant challenges to interpreting the cryo-EM/ET data using a single consensus model. Structural heterogeneity arises from multiple factors, including the inherent flexibility of macromolecules^9,10^, the presence of transient functional states, and the coexistence of different macromolecular species. These variations lead to substantial differences in the 3D structures of individual particles. The problem is further exacerbated in cryo-ET, where macromolecular complexes are visualized *in situ* and displays a much wider range of compositions and conformations. Addressing structural heterogeneity is thus a critical challenge in structural biology, as it is essential for discerning and segregating distinct conformations or species within samples. Traditional methods for resolving structural heterogeneity in cryo-EM/ET datasets often rely on repeated cycles of computationally intensive classification and particle subtraction procedures^7,11–15^. These approaches are not only time-consuming but also require extensive human intervention, limiting their applicability for high-throughput cryo-EM/ET studies. Moreover, these methods can only resolve a finite number of discrete states, making them inadequate for capturing continuous structural changes, which are crucial for understanding the functions of macromolecular complexes.

In recent years, deep learning has shown great promises in addressing complex problems across a wide range of fields owing to its exceptional modelling capacity^16^. In the context of cryo-EM data analysis, deep learning methods hold the potential to automate structural heterogeneity analysis. Several deep learning approaches to resolve 3D structural variations have been proposed. One class of methods employ end-to-end neural networks to translate 2D cryo-EM images into 3D volumes using results from traditional alignment algorithms. Examples include cryoDRGN^17^ and OPUS-DSD^18^, both of which capture structural variations using 3D volumes. While cryoDRGN represents the Fourier transform of the 3D volume with neural networks, OPUS-DSD directly uses 3D convolutional neural network (3DCNN) to represent 3D volume. Both methods have demonstrate the ability to resolve discrete and continuous structural heterogeneities^19^. Other methods, such as Multi-CryoGAN^20^, e2gmm^21^, and 3DFlex^22^, adopts different strategies. Multi-CryoGAN is a method based on generative adversarial network^23^. E2gmm uses a set of gaussians to model the 3D structure, while 3DFlex assumes each particle is derived from a common structure with variations modeled by a deformation field parameterized by multi-layer perceptron (MLP)^24^. For cryo-ET datasets, recent deep learning methods, such tomoDRGN^25^, and cryoDRGN-ET^26^ have extended the neural network architecture of cryoDRGN, leveraging high-resolution subtomogram averaging result to resolve structural heterogeneity in subtomograms.

Despite progress, key limitations persist. Methods like OPUS-DSD^18^ represents compositional and dynamic variations in a single latent space, reducing the interpretability of its result. Another limitation of OPUS-DSD lies in its inefficient dynamics modelling. Representing large-scale dynamics (e.g., subunit rigid-body motions) via voxel-level changes in 3D volumes incur substantial voxel changes, which is parametrically inefficient. Moreover, existing tools lack unification across cryo-EM/ET, limiting cross-modality insights.

Macromolecular dynamics are inherently hierarchical, dominated by low-frequency, large-scale normal modes (e.g., rigid-body subunit movements)^27^. Inspired by the multi-body refinement of RELION for cryo-EM data^28^ and the multi-group TLS method in X-ray crystallographic refinement^29^, parameterizing dynamics via subunit rigid-body transformations offers a low-dimensional, physically interpretable solution. Similarly, 3DFlex in cryoSAPRC^22^, and Zernike/spherical harmonic-based approaches^30,31^ demonstrate the potential of neural networks and mathematical basis functions for continuous deformation modeling.

We present OPUS-DSD2, a next-generation framework unifying structural heterogeneity analysis for cryo-EM/ET. Integrating concepts from Spatial Transformer^32^ and Neural Volumes^33^, OPUS-DSD2 employs MLPs to model the parameters of subunit-specific rigid-body dynamics, enabling efficient large-scale motion representation. By assuming the dynamics to be independent of macromolecular composition, OPUS-DSD2 encodes dynamics into a separate latent space, facilitating the disentanglement of composition changes and dynamics. Besides, OPUS-DSD2 employs a shared 3DCNN architecture to process both cryo-EM and cryo-ET data. It also adopts a unified 3D contrast transfer function (3D CTF)^4^ model to bridge imaging modalities, which is detailed in the image formation model subsection in **Methods**. In the **Results** section, we provide an overview of our approach, and delve into the architecture of OPUS-DSD2 for learning latent spaces pertaining to composition and dynamics. Systematic testing of OPUS-DSD2 on real cryo-EM/ET datasets showcases high quality structural heterogeneity resolving result for macromolecules with biological insights for their functions.

## Results

### Network Architecture

To reveal the structural differences in space for a macromolecule, particles with different poses must be aligned to the canonical orientation of a model. OPUS-DSD2 leverages pose parameters derived from traditional alignment algorithms. For cryo-EM, the consensus refinement^7,34^ determines the translation and orientation of images or subtomograms relative to the reference model. For cryo-ET, template matching^8^ determines the location and orientation of a specific macromolecule inside a tomogram, while subtomogram averaging^4^ determines the translation and orientation of subtomograms relative to the reference model.

OPUS-DSD2 resolves structural heterogeneity by reconstructing the 3D structure for each 2D cryo-EM image or 3D cryo-ET subtomogram. The reconstruction of a 3D structure by the neural architecture of OPUS-DSD2 follows the steps outlined below (**Fig.1a**). The 3D subtomogram serves as the common input format for OPUS-DSD2. For cryo-EM data, a 2D image is tiled along the *z* axis to create a 3D subtomogram before being supplied into the neural network. The 3D subtomogram is then aligned to the canonical orientation of a consensus model by rotating it inversely based on its estimated pose (**Fig.1a**). The encoder processes the aligned 3D subtomogram and estimates the distributions of two latent codes. The latent code for the composition decoder, referred to as the composition latent code *z_c_*, has *n*-dimensions, while the latent code for the dynamics decoder, referred to as dynamics latent code *z_d_*, has *m*-dimensions. Both latent codes are assumed to follow Gaussian distributions, with means *z* ∈ ℝ*^n^* and standard variations σ ∈ ℝ*^n^*, which are estimated by the encoder^35^. The two latent codes are sampled from their respective distributions and fed into corresponding decoders. The sampled composition latent code is supplied to the composition decoder to generate a 3D volume (**Fig.1a**), which is represented by a discrete grid of voxels *V*(***x***), where ***x*** ∈ ℝ^3^ denotes a grid point in 3D space. Meanwhile, the sampled dynamics latent code is supplied to the dynamics decoder to generate a deformation field (**Fig.1a**), which is represented by 3D vectors on a discrete grid, i.e., ***g***(***x***): ℝ^3^ → ℝ^3^. Each vector at a grid point ***x*** defines the source voxel contributing to the output at position ***x***. The deformation field is constructed using the rigid-body displacement of each subunit with the parameters predicted by the dynamics decoder, as described in the **Deformation field** subsection in **Methods**. Using a spatial transformer^32^ network, ***g***(***x***) warps the 3D volume *V*(***x***) to produce a deformed volume. The deformed volume is subsequently transformed into a reconstruction with estimated pose and CTF parameters following the differentiable subtomogram formation model. The neural architecture of OPUS-DSD2 is trained end-to-end by reconstructing each input image or subtomogram and minimizing the squared reconstruction error.

**Figure 1.**
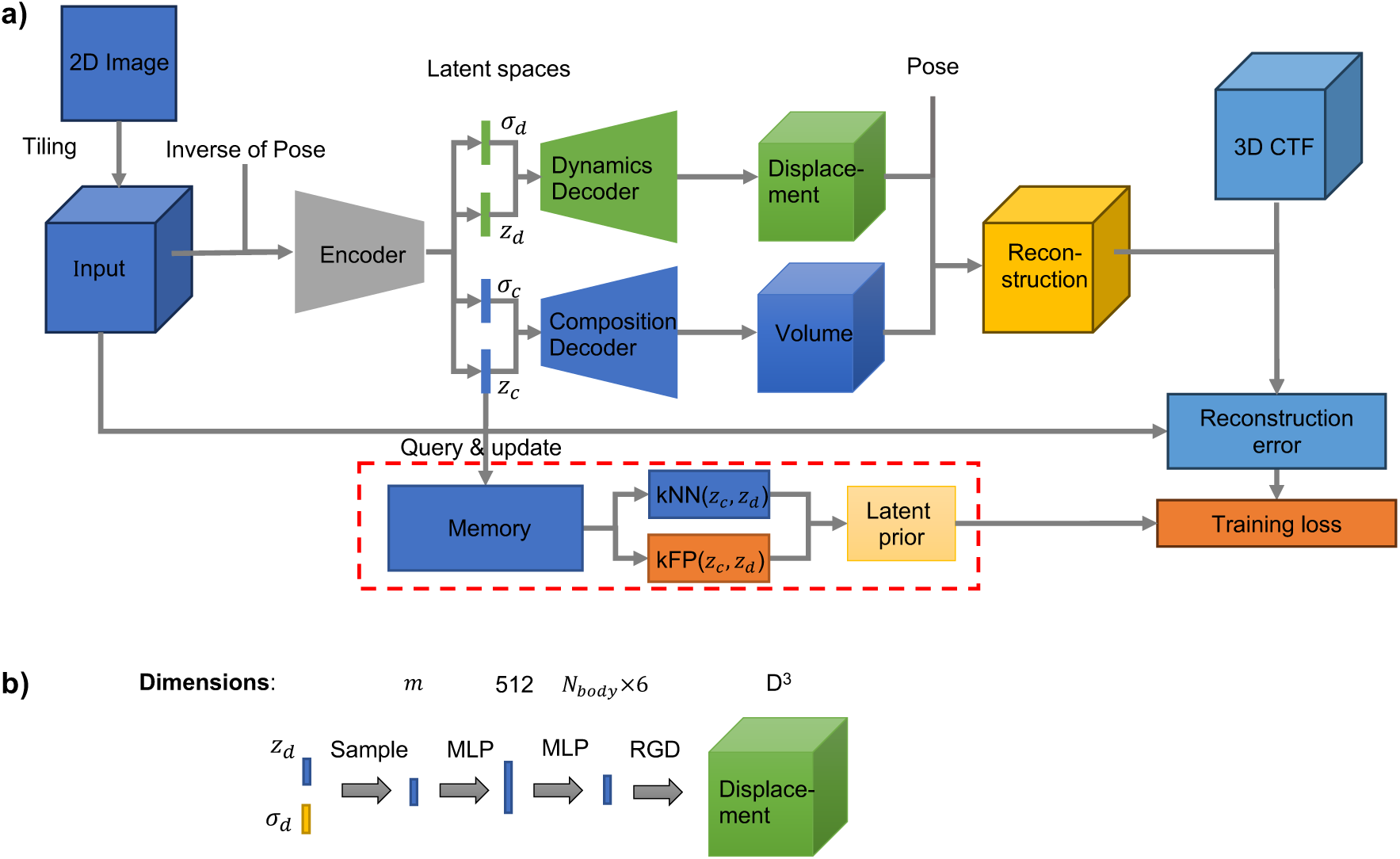
Architecture of OPUS-DSD2. **a)**. **Overall architecture.** Tiling refers to tiling the 2D image along *z* axis, which is only applied to cryo-EM image. Pose refers to the projection direction of input with respect to the consensus model. Inverse of Pose refers to the inverse of projection direction of input with respect to the consensus model. 3D CTF refers to 3D contrast transfer function. *z_d_* is the mean vector of composition latent code. σ_c_ is the standard deviation vector of composition latent code. *z_d_* is the mean vector of dynamics latent code. σ*_d_* is the standard deviation vector of dynamics latent code. kNN(*z_c_*, *z_d_*) refers to the K nearest neighbor of the concatenation of *z_d_* and *z_d_*. kFP(*z_c_*, *z_d_*) refers to the K farthest points of the concatenation of *z*_!_and *z_d_*. For better visualization, the link between the dynamics latent code and the memory is omitted. For simplicity, the standard gaussian prior for latent distribution as well as the smoothness and sparseness priors for the 3D volume are omitted in this chart. **b)**. **Architecture of the dynamics decoder of OPUS-DSD2.** This diagram shows the dynamics decoder that translates the latent encoding *z_d_* into a deformation field. MLP denotes the multi-layer perceptron. RGD denotes the rigid-body dynamics model that transforms the predicted rigid-body movement parameters into the 3D deformation field. All MLPs except last one are with LeakyReLU nonlinearities with negative slope 0.2.

During training, to encourage the smoothness of latent spaces, the composition and dynamics latent codes are regularized by the structural disentanglement prior from OPUS-DSD^18^, as well as the Kullback-Leibler (KL) divergence with respect to (w.r.t) standard gaussian, as in variational autoencoder (VAE)^35,36^. The composition decoder utilizes the 3D convolutional network from OPUS-DSD^18^. The dynamics decoder consists of a series of MLPs and nonlinearities (**Fig.1b**). The dynamics latent code is transformed into a 512-dimensional representation by an MLP followed by a LeakyReLU nonlinearity. The 512-dimensional representation is projected to a *N*_body_ × 6-dimensional vector by another MLP, where *N*_body_ denotes the number of rigid bodies defined for the macromolecule. This *N*_body’_× 6-dimensional vector is then reshaped into *N*_body_ vectors, each of 6 dimensions, which define the rotation and translation parameters for each subunit’s rigid-body dynamics. The first three dimensions of the 6-dimensional output define the rotation using a quaternion representation, while the last three dimensions defines the translation of the subunit. The construction of the deformation filed using the rotation and translation parameters is detailed in the **Deformation field** subsection in **Methods**.

### Pre-catalytic spliceosome

We tested the performance of OPUS-DSD2 in structural heterogeneity analysis for cryo-EM data on the pre-catalytic spliceosome (EMPIAR-10180)^37^. We trained a 16-dimensional latent variable model, where 12 dimensions are used for composition latent space and the remaining 4 dimensions are kept for dynamics latent space. The consensus refinement result for model training is from EMPIAR-10180^38^. The rigid-body dynamics are defined according to the subunit masks deposited in EMPIAR-10180^28^. We first compared the reconstruction qualities of the volumes from OPUS-DSD2 to the volumes from OPUS-DSD which was trained with 12-dimensional latent space. Distinct conformations are revealed by principal component analysis (PCA) on the composition latent space. At the positive end of the first principal component (PC1) of OPUS-DSD2 and PC1 of OPUS-DSD, the reconstructed template volume corresponds to an ‘open’ conformation where the SF3b subunit stays on the core body, while the reconstructed volume at the negative ends of PCs corresponds to a ‘closed’ conformation where SF3b subunit folds onto the back of core body. **Fig.2a** shows the ‘open’ conformation reconstructed by OPUS-DSD2 and OPUS-DSD. Contoured at the same level, the reconstruction of OPUS-DSD2 shows clearly separated helices in front view in the top panel. In contrast, the reconstruction of OPUS-DSD presents blurred densities at the same regions (**Fig.2a**). The difference become more evident in the enlarged views inside the boxed regions. The reconstruction from OPUS-DSD2 shows more resolved structural folds in the SF3b subunit (**Fig.2a**). **Fig.2b** shows the ‘closed’ conformation reconstructed by OPUS-DSD2 and OPUS-DSD. Similarly, in the front view, the reconstruction from OPUS-DSD2 shows clearer and more continuous structures at both Core and Helicase subunits (**Fig.2b**). In the rear view, the improved structural features in the reconstruction of OPUS-DSD2 compared to the reconstruction from OPUS-DSD are noticeable (**Fig.2b**). Traversal along different PCs in the composition latent space of OPUS-DSD2 corresponding different changes of components in structures. We compared the traversals along PC3 and PC5 side by side, and observed emergences of two different structural elements between the Foot and the Helicase (**Supplementary Video 1**).

**Figure 2.**
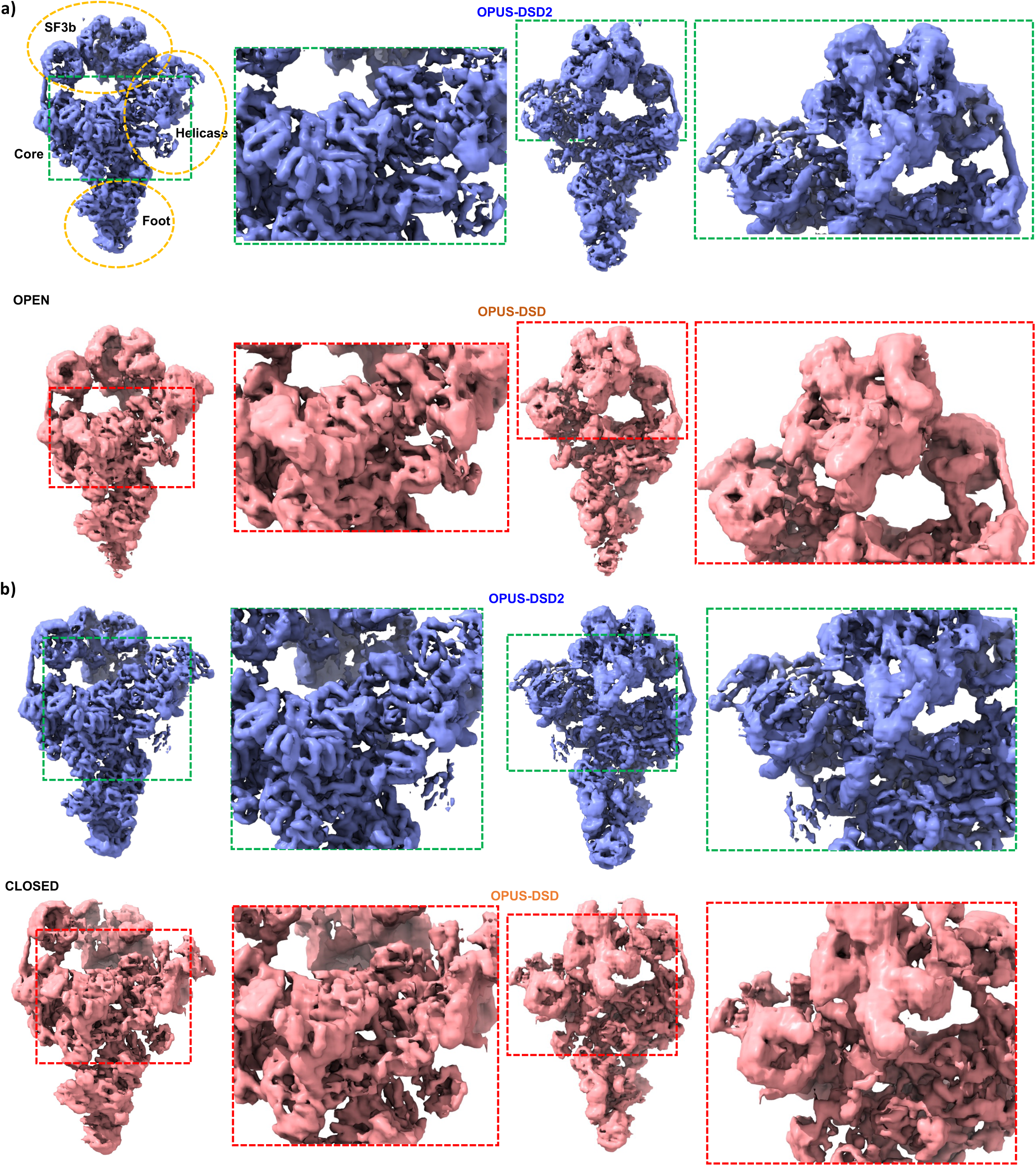
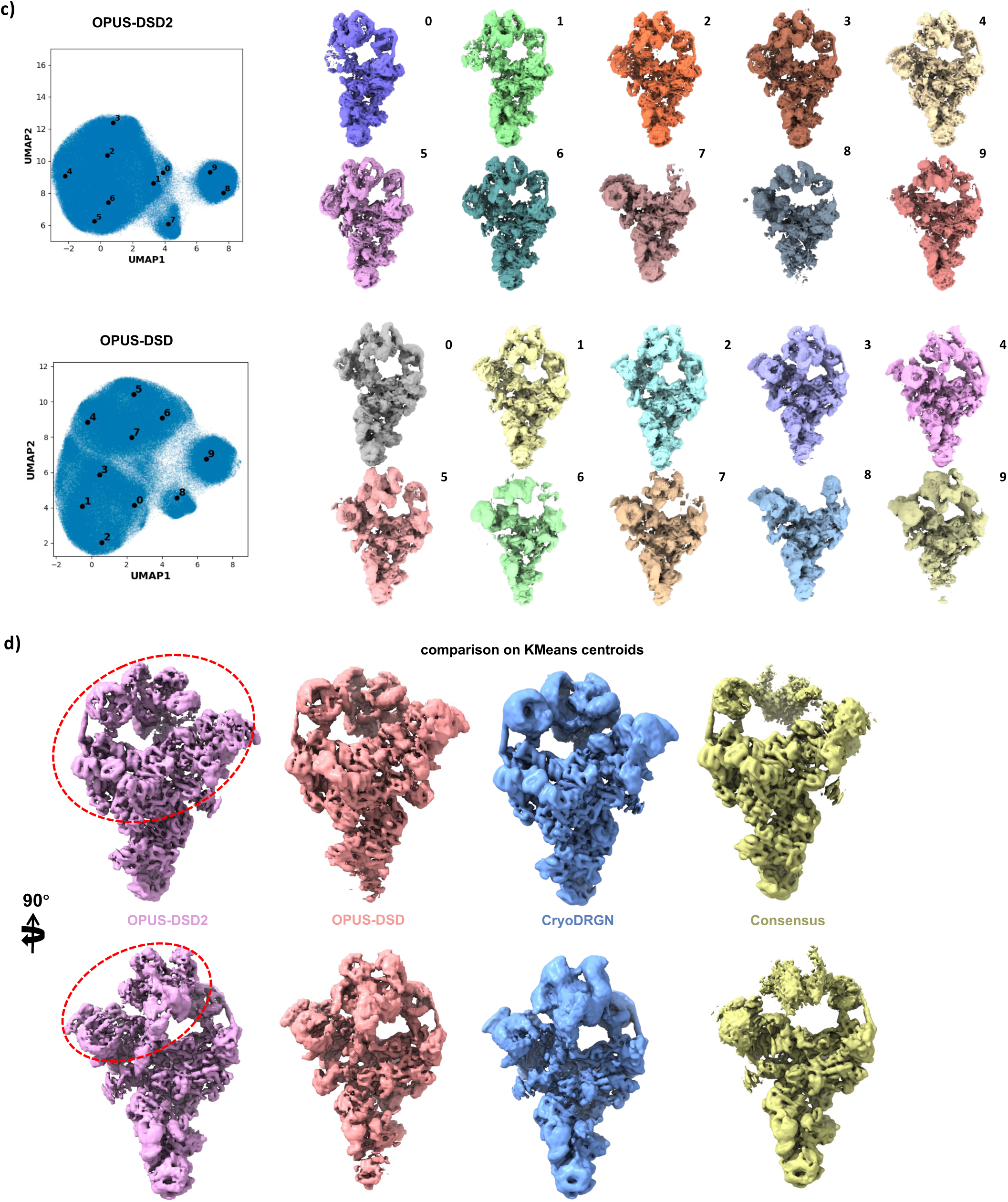
Heterogeneity analysis on the pre-catalytic spliceosome (EMPIAR-10180). **a)**. Open conformations of spliceosome generated along the PC1 of latent space from OPUS-DSD2 and the PC1 of latent space from OPUS-DSD. The open conformations are generated at PCs with amplitude -1.1. **b)**. Closed conformations of spliceosome generated along the PC1 of the latent space from OPUS-DSD2 and the PC1 of latent space from OPUS-DSD. The closed conformations are generated at PCs with amplitude -1.1. The regions with large differences in the qualities of densities are marked and highlighted by dashed boxes. All maps are contoured at the same level. **c)**. UMAP^58^ visualizations of the 12-dimensional latent spaces of all particles encoded by OPUS-DSD2 and OPUS-DSD. Solid black dot represents the cluster center for labelled class. The corresponding reconstructions at the cluster centers are shown as well. **d)**. Closed conformations of spliceosome generated at the centroids of KMeans clusters in latent spaces learned by OPUS-DSD2, OPUS-DSD and CryoDRGN.

We also compared the reconstruction qualities of different methods on the KMeans^39^ centroids (Fig.2c and **Fig.2d**). The latent spaces of the results of both methods are clustered into 10 classes using KMeans algorithms (**Fig.2c**). The reconstructions from OPUS-DSD2 are of higher resolutions than the reconstruction from OPUS-DSD. Similar reconstructions from the results of different methods are selected for close-up comparison (**Fig.2d**). In regions marked by red dashed boxes, the reconstruction from OPUS-DSD2 presents structural features with higher resolution than reconstructions from other methods (**Fig.2d**).

It’s worth noting that the qualities of reconstructions differ at KMeans centers and the ends of PCs. This is caused by that KMeans and PCA are different ways to sample the latent space. The centers from KMeans clustering are in dense regions, while the points on principal components are manually set. Therefore, the reconstructions at centers from KMeans clustering are generally of higher resolution, while the reconstructions corresponding to points at two ends of PCs are of lower resolution owing to the sparseness of the latent encodings around them.

Another important functionality of OPUS-DSD2 is reconstructing dynamics explicitly by traversing along the PCs of dynamics latent space (DPC). We performed PCA on the dynamics latent space. Class 2 from KMeans clustering in the composition latent space was chosen as the template conformation to be deformed. Traversing the first PC reveals the concerted folding of SF3b and Helicase towards Core (**Supplementary Video 2**), and traversing the second PC reveals the folding of Helicase towards Core, while the SF3b rotating around its center (**Supplementary Video 3**). The dynamics reconstructed by OPUS-DSD2 shows continuous deformation of densities near boundaries between subunits, which is in sharp contrast to the dynamics reconstructed by Multi-body refinement in RELION^28^, where densities near boundaries between subunits are broken.

### Plasmodium falciparum 80S ribosome

We tested the performance of OPUS-DSD2 in structural heterogeneity for cryo-EM data on *Plasmodium falciparum* 80S (Pf80S) ribosome (EMPIAR-10028)^40^. We trained a 16-dimensional latent variable model, where composition latent space is of 12 dimensions and dynamics latent space is of 4 dimensions. We followed the subunit segmentation scheme in the multibody refinement of Pf80S^28^, except that the Head and SSU subunits are merged as a single subunit in this work. Hence, the deformation field in our analysis is comprised of movements of two subunits. We first compared the reconstruction qualities of volumes from the 12-dimensional model of OPUS-DSD2 to volumes from the model of OPUS-DSD with the same number of dimensions. PCA is performed on both latent spaces. Reconstructions at points on the PC3 from OPUS-DSD2 and the PC1 from OPUS-DSD are compared. We denoted the conformations at the positive end of PCs, with magnitude being 1, as ‘POS’. At the same contour level, the reconstructions from both methods have similar compositions at this point (**Fig.3a**). However, in both viewing directions, the reconstruction from the result of OPUS-DSD2 shows higher resolution structures as exemplified by the enlarged views inside dashed boxes (**Fig.3a**). For example, the separation between parallel helices is more evident in the reconstruction from OPUS-DSD2 (**Fig.3a**). Next, we denoted the conformations at the center of PCs, with magnitude being 0.1, as ‘NEU’ (**Fig.3b**). The reconstruction by OPUS-DSD2 still demonstrates superior qualities (**Fig.3b**). By traversing along different PCs of the latent space from OPUS-DSD2, different patterns of compositional changes were revealed (**Supplementary Video 4**). The PC1 mainly encodes the compositional changes within the Head in SSU, that is, several RNAs go from presence to missing during the twisting of Head (**Supplementary Video 4**). The PC3 mainly encodes the compositional changes of an RNA strand connecting SSU and LSU. The RNA strand slides from SSU to LSU when traversing along the PC3 (**Supplementary Video 4**).

**Figure 3.**
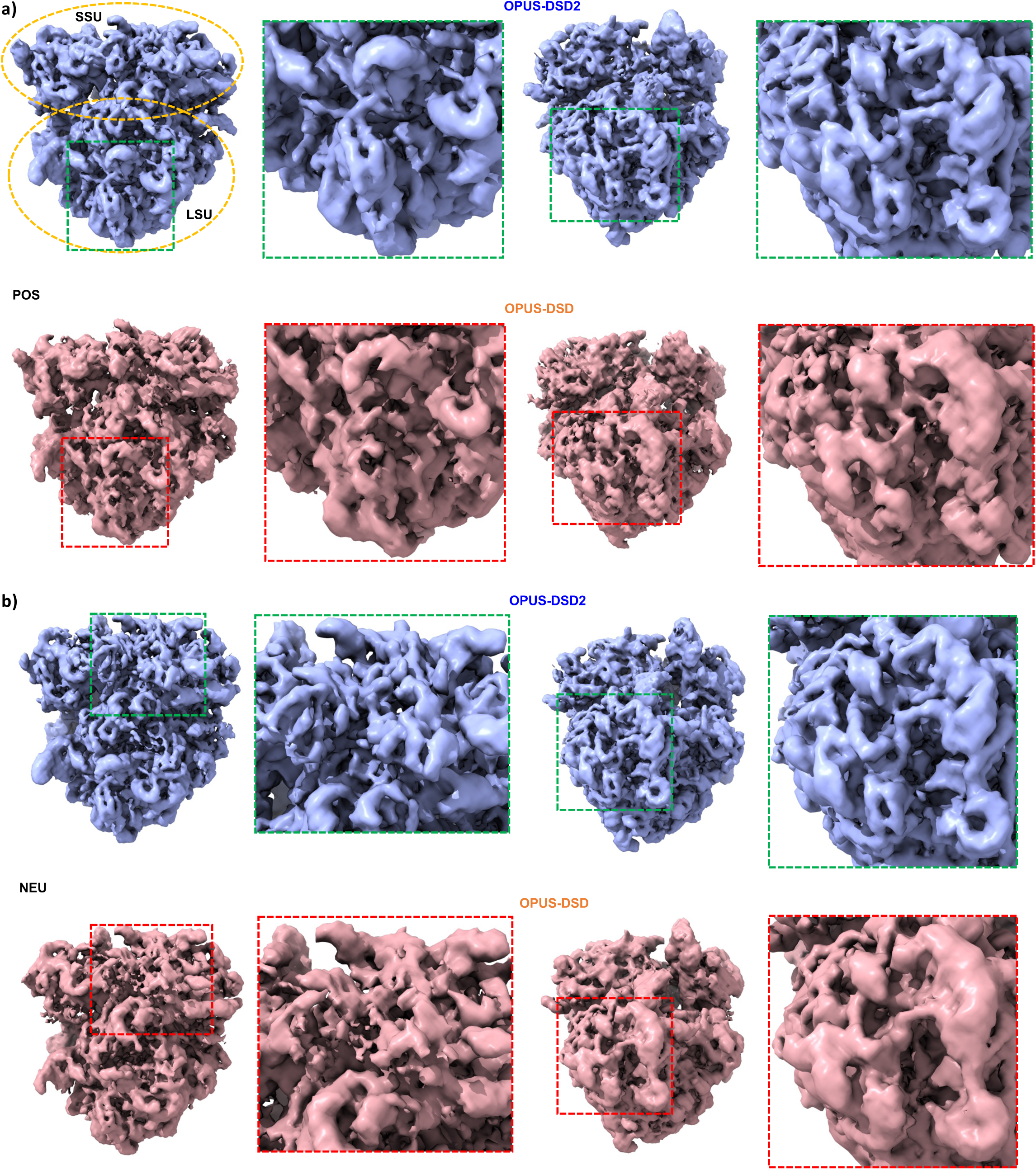
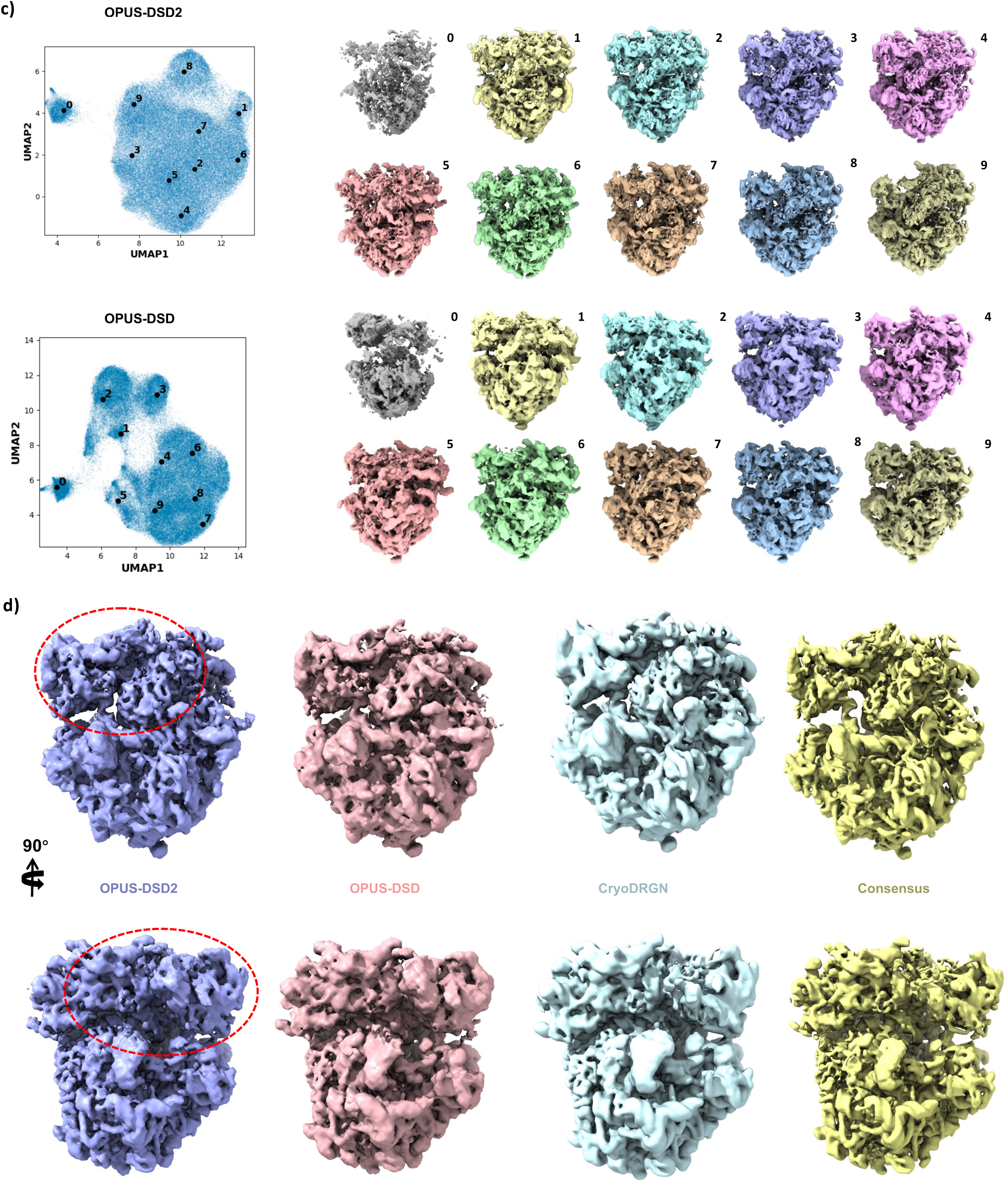
Heterogeneity analysis on *Plasmodium falciparum* 80S ribosome (EMPIAR-10028). **a)**. POS conformations of Pf80S ribosome generated along the PC3 of latent space learned by OPUS-DSD2 and the PC1 of latent space learned by OPUS-DSD. The POS conformations are generated at PCs with amplitude 1. **b)**. NEU conformations of Pf80S ribosome generated along the PC3 of latent space learned by OPUS-DSD2 and the PC1 of latent space learned by OPUS-DSD. The NEU conformations are generated at PCs with amplitude 0.1. The regions with large differences in the qualities of densities are marked and highlighted by dashed boxes. All maps are contoured at the same level. **c)**. Conformations of Pf80S ribosome generated at the centroids of KMeans clusters in latent spaces learned by OPUS-DSD2, OPUS-DSD and CryoDRGN. **d)**. UMAP visualizations of the 12-dimensional latent spaces of all particles encoded by OPUS-DSD2 and OPUS-DSD. Solid black dot represents the cluster center for labelled class. The corresponding reconstructions at the cluster centers are shown as well.

We compared the qualities of reconstructions on centroids from KMeans clustering in latent spaces of both methods. Both latent spaces are clustered into 10 classes using KMeans algorithms. Volumes are generated by supplying the centroids into the decoders of OPUS-DSD2 and OPUS-DSD, and are visualized at the same contour level in **Fig.3c**. The reconstructions by OPUS-DSD2 consistently present more structural details. In the selected reconstructions on KMeans centroids, the reconstruction by OPUS-DSD2 shows more separated structural elements. (**Fig.3d**).

We performed PCA on the dynamics latent space, and visualized the learned multi-body dynamics by traversing along PCs. Class 4 from KMeans clustering was chosen as the starting conformation to be deformed. The relative rotation between the LSU and SSU can be revealed by traversing the second PC of the dynamics latent space (**Supplementary Video 5**).

### SARS-CoV-2 spike protein trimer

We tested the performance of OPUS-DSD2 in structural heterogeneity for cryo-EM data on a SARS-CoV-2 spike (S) protein trimer (EMPIAR-10492)^41^. S protein trimer is comprised of two regions, S1 and S2 (Fig.4a). S1 region can be further divided into two types of subunits, N-terminal domain (NTD) and C-terminal domain (CTD) (**Fig.4a**). S1 CTDs serve as the receptor-binding domain, and is found to present two different compositions in this dataset by previous study^41^, which have three closed CTDs and two closed CTDs, respectively. Given the *C*_3_symmetry inherent to the trimer, we applied symmetry expansion to the pose parameters of images deposited in EMPIAR-10492 using RELION 3.0.8. Following this symmetry expansion, OPUS-DSD2 was trained on images with the assumption that their projection angles correspond to one of the three equivalent angles satisfying *C*_3_symmetry. We trained a 16-dimensional latent variable model, where the composition latent space is of 12 dimensions and the dynamics latent space is of 4 dimensions. S trimer is segmented to 4 subunits as it is shown in **Fig.4a**. KMeans clustering was performed on the composition latent space to classify particles into 16 classes (**Fig.4b**). The reconstructions produced by the decoder of OPUS-DSD2 at cluster centers reveal different structures of S trimer, some of which show three closed CTDs while some have one CTD missing (**Fig.4c**).

**Figure 4.**
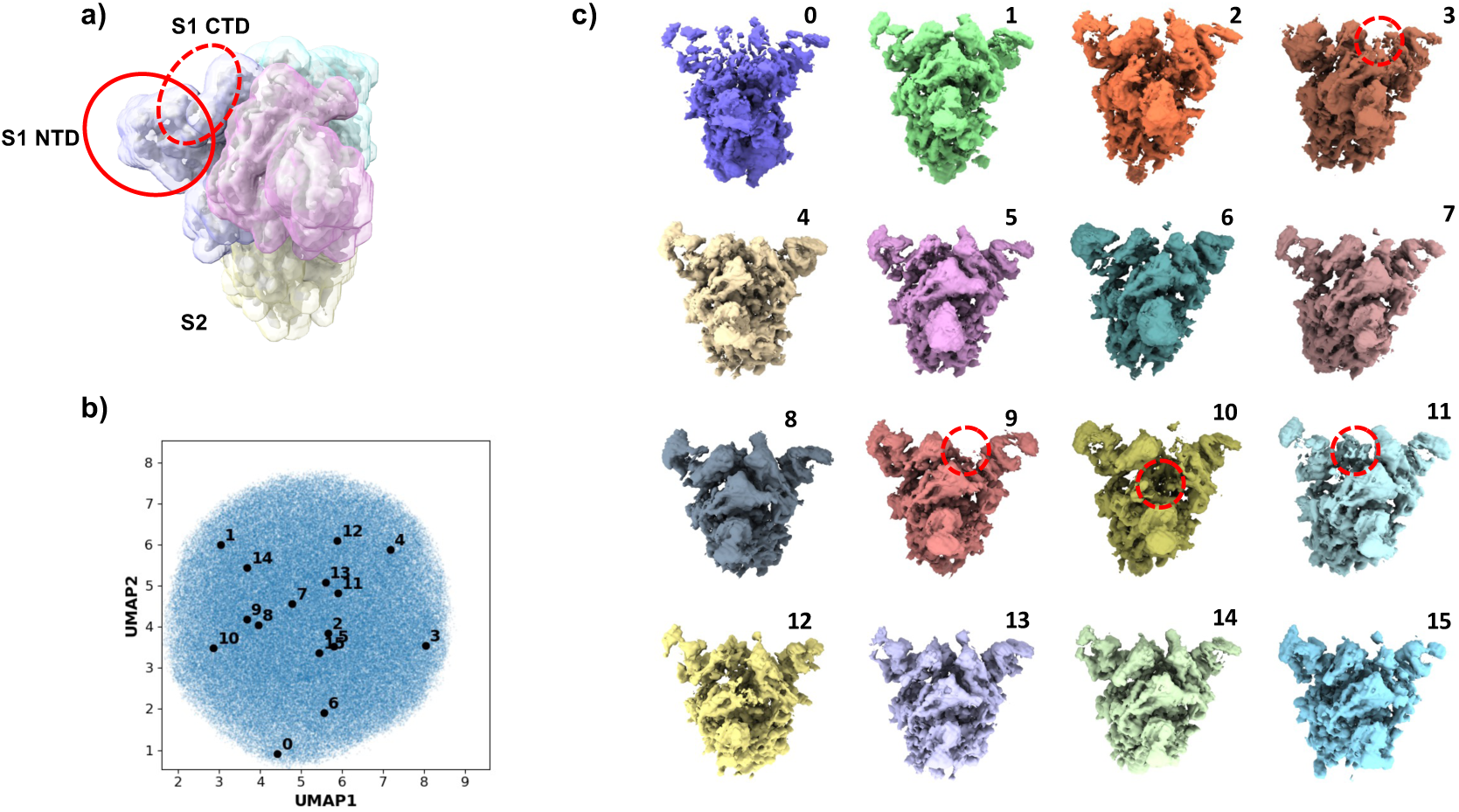
Heterogeneity analysis on SARS-CoV-2 spike protein trimer (EMPIAR-10492). **a)**. The segmentation of SARS-Cov-2 spike protein trimer. **b)**. UMAP visualization of the 12-dimensional composition latent space learned by OPUS-DSD2. Solid black dot represents the cluster center for labelled class. **c)**. Structures of S trimer generated by the decoder of OPUS-DSD2 at the centroids of KMeans clusters in the composition latent space. Red dashed circles mark the missing S1 CTD in the corresponding structures.

The dynamics and composition latent space encode dynamics in different scales. Two orthogonal movement modes for S1 subunits can be found in the dynamics latent space. Traversing the first principal component (DPC1) of the dynamics latent space shows a symmetric dynamics where all monomers are moving towards the center synchronously, which leads to the open and close of S trimer (**Supplementary Video 6**). Traversing the second principal component (DPC2) of the dynamics latent space reveals an asymmetric dynamics where the upper S1 subunit is moving in the opposite direction relative to the center compared to other subunits (**Supplementary Video 6**). The dynamics within S1 NTD are embedded in the composition latent space. For example, traversing the PC3 of composition latent space corresponds to the open and close of NTD (**Supplementary Video 7**).

### *S. Pombe* 80S Ribosome

We tested the performance of OPUS-DSD2 in the structural heterogeneity analysis of cryo-ET data on *Schizosaccharomyces pombe* (*S. Pombe*) 80S ribosome *in situ*. The publicly available tomograms of wild-type *S. pombe* from DeePiCt’s work^42^ (EMPIAR-10988) were used for this analysis. A total of 50,000 particles corresponding to 80S ribosomes were picked by template matching in pyTOM^43^ using the published *S. cerevisiae* 80S ribosome map^4^ (EMDB accession code: EMD-3228) as the template. Initially, non-ribosome particles were filtered from the template matching result by training OPUS-TOMO^44^ on subtomograms and their orientations from the template matching result directly. The resulting latent space was clustered into 20 classes, with class 12∼17 consisting of 17,215 subtomograms exhibiting densities for 80S ribosome (**Fig.5a∼b**). These classes were selected for further structural heterogeneity analysis in OPUS-DSD2. The rigid-body dynamics for 80S ribosome was defined as the relative displacement between 40S and 60S subunits. We then trained OPUS-DSD2 using the 17,215 subtomograms with their orientations from the template matching result directly. The composition latent space is set to 12 dimensions and the dynamics latent space is set to 4 dimensions.

**Figure 5.**
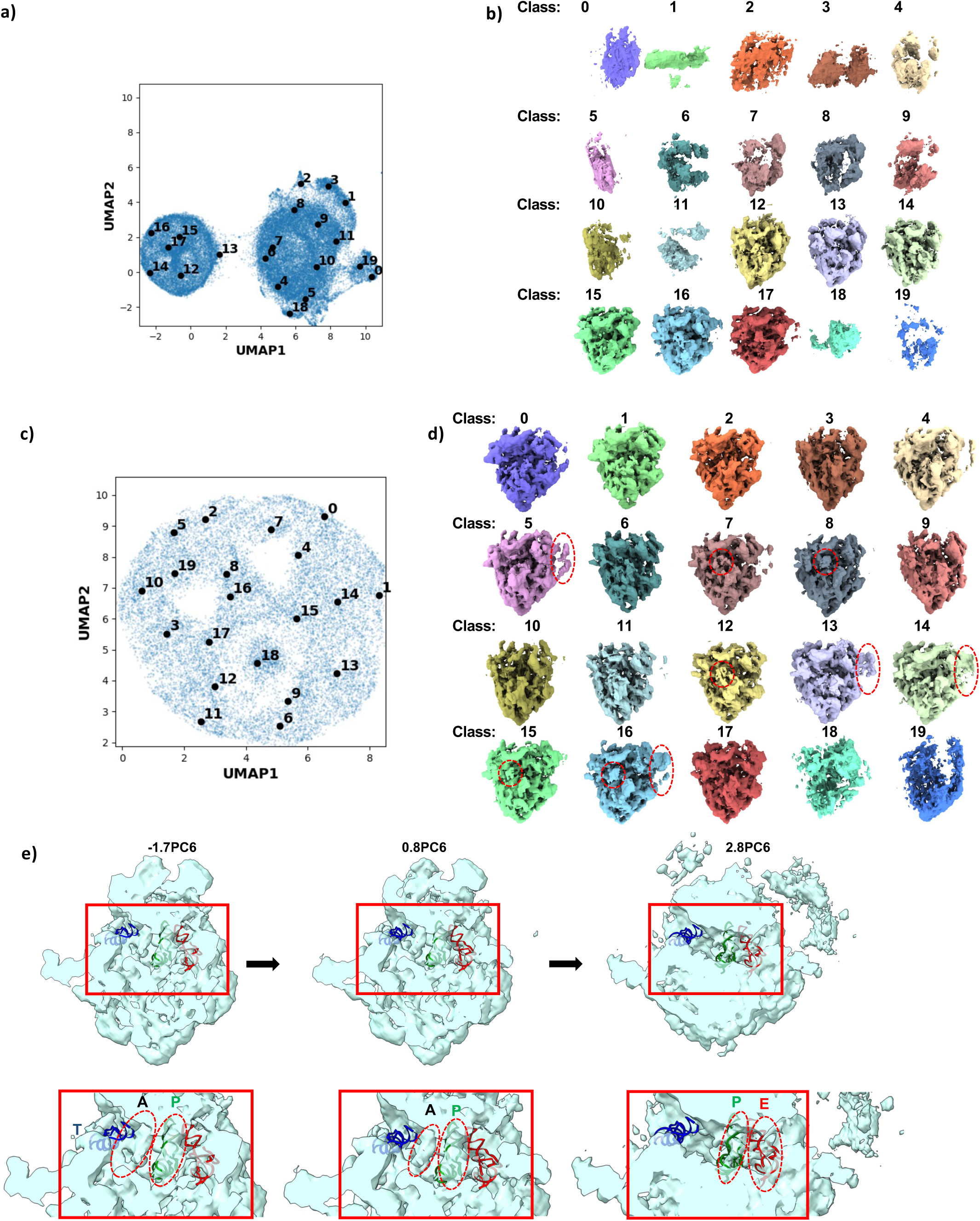
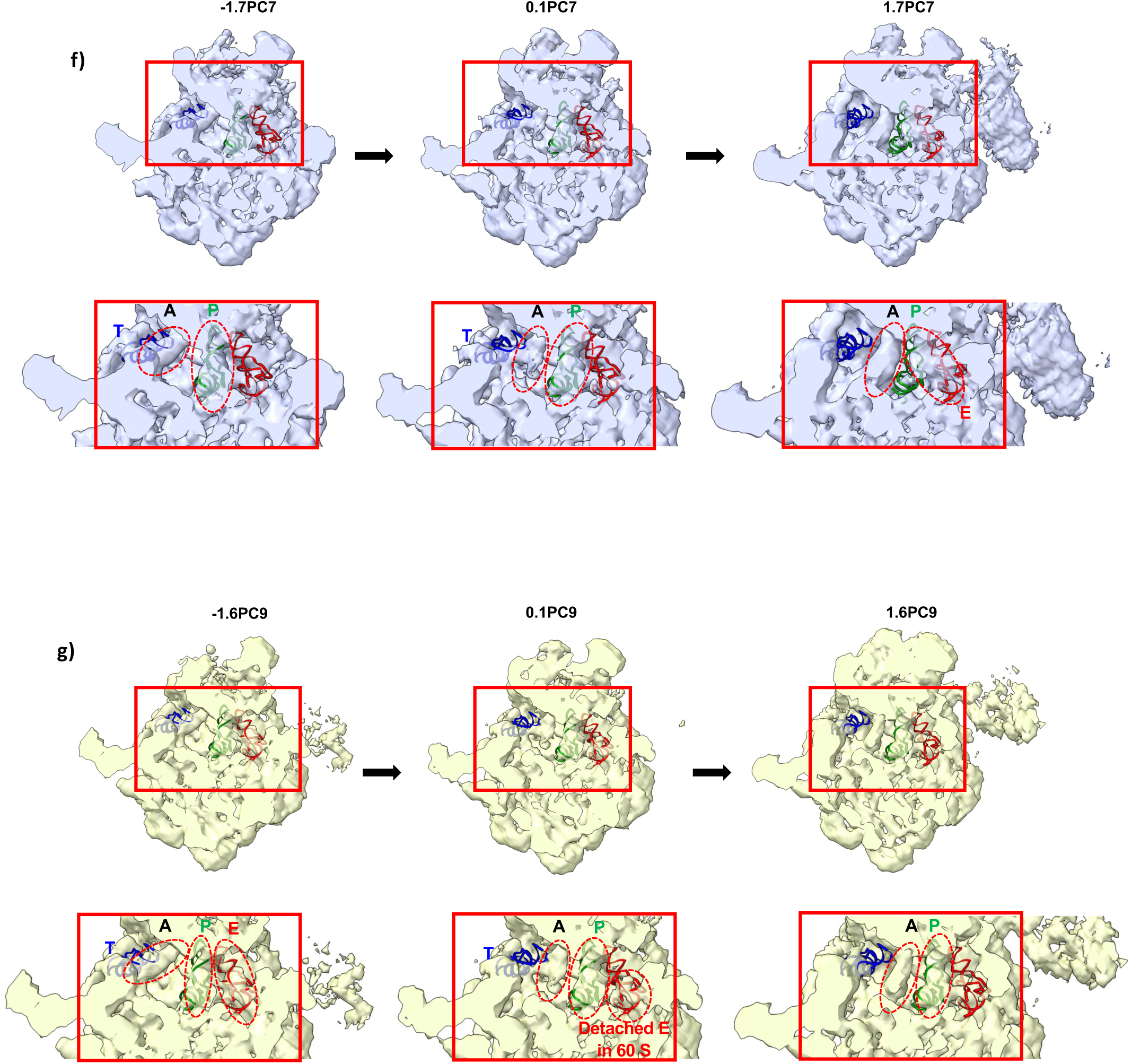
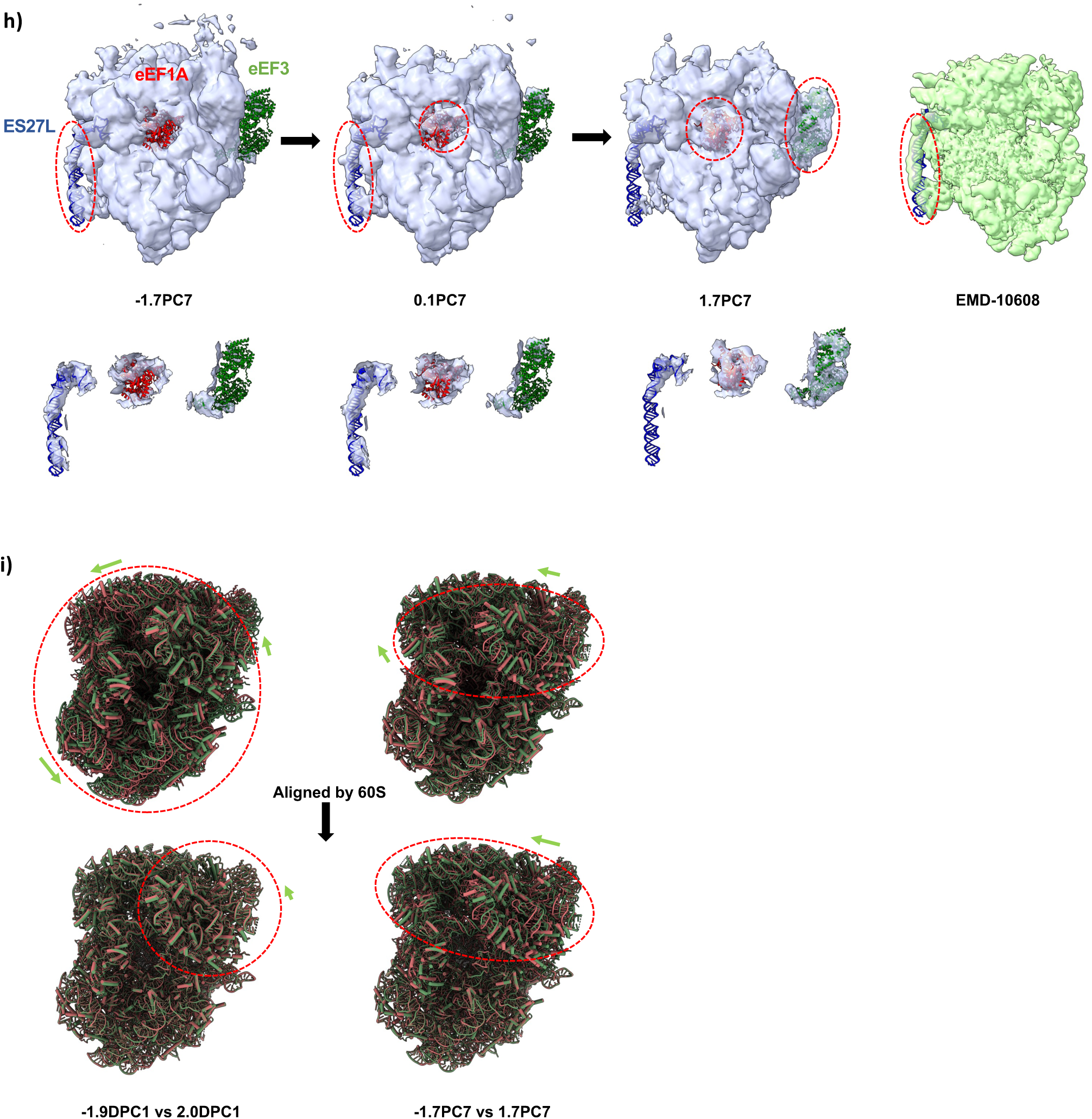
Heterogeneity analysis on *S. pombe* 80S ribosome (EMPIAR-10988). **a)**. UMAP visualization of the 12-dimensional composition latent space learned by OPUS-TOMO. Solid black dot represents the cluster center for labelled class. **b)**. Structures of *S. pombe* 80S ribosome generated by the decoder of OPUS-TOMO at the centroids of KMeans clusters in the composition latent space. **c)**. UMAP visualization of the 12-dimensional composition latent space learned by OPUS-DSD2. Solid black dot represents the cluster center for labelled class. **d)**. Structures of *S. pombe* 80S ribosome generated by the decoder of OPUS-DSD2 at the centroids of KMeans clusters in the composition latent space. Additional cofactors, eEF1A and eEF3 are marked by red dashed ellipses. **e)**. The transient states for tRNA translocation reconstructed by the Decoder of OPUS-DSD2 using latent codes at different locations along PC6. The locations of latent codes are specified as xPCn. The sites occupied by tRNAs are marked by red dashed ellipses and annotated with corresponding labels. The atomic structure of *S. c* 80S ribosome (PDB code: 9BDP) with A/T-, P- and E-site tRNAs was fitted into these density maps. The A/T-site tRNA was colored in blue, the P-site tRNA was colored in green, and the E-site tRNA was colored in red. The A/T-site consists of A-site at 40S subunit and T-site at 60S subunit. **f)**. The transient states for tRNA translocation reconstructed by the Decoder of OPUS-DSD2 using latent codes at different locations along PC7. The sites occupied by tRNAs are marked by red dashed ellipses and annotated with corresponding labels. The A/T-site consists of A-site at 40S subunit and T-site at 60S subunit, the hybrid P/E-site consists of P-site at 40S subunit and E-site at 60S subunit. **g)**. The transient states for tRNA translocation reconstructed by the Decoder of OPUS-DSD2 using latent codes at different locations along PC9 in composition latent space. The sites occupied by tRNAs are marked by red dashed ellipses and annotated with corresponding labels. The A/T-site consists of A-site at 40S subunit and T-site at 60S subunit. **h)**. Occupancy changes of different cofactors in density maps reconstructed by the Decoder of OPUS-DSD2 using latent codes at different locations along PC9 in composition latent space. The sites occupied by additional ES27L RNA and elongation cofactors, eEF1A and eEF3, are marked by red dashed ellipses and annotated with corresponding labels. **i)**. Relative displacements between 40S subunit and 60S subunit in density maps reconstructed by the Decoder of OPUS-DSD2 using latent codes at different locations along DPC1 in dynamics latent space and PC7 in composition latent space. The structures fitted with the density maps reconstructed by the Decoder of OPUS-DSD2 at the -1.9DPC1 and -1.7PC7 are colored in green, and the structures fitted with the density maps reconstructed by the Decoder of OPUS-DSD2 at 2.0DPC1 and 1.7PC7 are colored in red. The green arrow denotes the direction of displacement from the green structure to red structure.

The composition latent space learned by OPUS-DSD2 was clustered into 20 classes. Classes 13 and 14 showed additional densities for eEF3, while classes 7∼8 and 16 displayed additional densities for eEF1A (**Fig.5c∼d**). The 80S ribosome *in situ* undergoes a constant translation elongation cycle to synthesize polypeptide from mRNA^45^. Specifically, the 80S ribosome has three tRNA binding sites, A, P and E sites. During elongation, these sites are sequentially occupied and released by tRNAs, facilitating the translation of mRNA to peptide chains. Impressively, the complex compositional and conformational changes underlying different stages of translation elongation cycle can be revealed by traversing the PCs of the composition latent space learned by OPUS-DSD2.

Using the atomic structure of *Saccharomyces cerevisiae* (*S. c*) 80S ribosome (PDB code: 9BDP)^46^ as reference, traversal of the PC6 reveals the translocation dynamics of tRNAs, starting from a state where the A-site is partially occupied and the P-site is fully occupied (-1.7PC6), transitioning through a state where both A- and P-site are occupied (0.8PC6), and culminating in a state where P- and E-site are occupied (2.8PC6) (**Fig.5e**). Similarly, traversal of PC7 shows the transition from a state with A/T tRNA and P-site being occupied, through a transient state with partial occupation of A-site and full occupation of P-site, ending in a state with A-site tRNA and hybrid P/E-site tRNA (**Fig.5f**). Moreover, traversal of PC9 demonstrates the transition from a state where A/T tRNA, P- and E-site are occupied, through a transient state where A-site is partially occupied, P-site is occupied and E-site tRNA is detached from 40S subunit, ending in a steate where A-site is occupied and E-site is fully released (**Fig.5g**). Traversal of PC7 can also reveal compositional changes related to translation elongation factors which bind with 80S ribosome and the additional ES27L RNA at exit tunnel (**Fig.5h**). The state at -1.7PC7 shows densities for the ES27L RNA at exit tunnel, and highly incomplete densities for cofactors eEF1A and eEF3. The state at 0.1PC7 shows densities for the ES27L RNA at exit tunnel and the cofactor eEF1A, while showing highly incomplete densities for cofactors eEF3. The state at 1.7PC7 shows no densities for the additional ES27L RNA at exit tunnel, and densities for cofactors eEF1A and eEF3.

To dissect dynamics of 80S ribosome, we analyzed the OPUS-DSD2-derived dynamics latent space (DPC1) and composition latent space (PC7). The atomic structure for *S. c* 80S ribosome (PDB code: 9BDP)^46^ was fitted into the density maps reconstructed by OPUS-DSD2 along DPC1 (-1.9DPC1 and 2.0DPC1) (**Fig.5i**). Both the 40S subunit and 60S subunit were fitted separately. The fitted structures revealed large-scale rigid-body displacements in both subunits (Fig.5i **upper left panel**). Alignment of these structures via the 60S subunit highlighted small residual motion in the 40S head domain (**Fig.5i lower left panel**). In contrast, traversal of PC7 (-1.7PC7 to 1.7PC7) showed minimal 60S movement but pronounced 40S rearrangements, indicating localized subunit dynamics (Fig.5i **right panel**). These results demonstrate that the dynamics latent space (DPC1) captures global ribosomal motions, likely arising from subtomogram alignment errors during templating matching, while the composition latent space (PC7) resolves localized conformational changes. The integration of dynamics model effectively corrects pose assignment inaccuracies, enhancing reconstruction quality and enabling resolution of ribosome dynamics at multiple scales.

### *S. Pombe* 26S Proteasome

We tested the performance of OPUS-DSD2 in the structural heterogeneity analysis for cryo-ET data on *Schizosaccharomyces pombe* (*S. Pombe*) 26S proteasome *in situ*, a macromolecular complex responsible for protein degradation in eukaryotic cells. The 26S proteasome consists of a 20S cylindrical core particle (CP) flanked by two 19S regulatory particles (RPs) at each end^15^. The RP consists of a ring-shaped AAA-ATPase heterohexamer (Rpt1-6) surrounded by 13 RP non-ATPases (Rpn1-3, 5-13, 15)^47^. The RPs serve to recruit ubiquitylated substrates, cleave off their polyubiquitin tags, and unfold and translate them into the CP for degradation^15^. The functions of RPs are carried out in coordination with deubiquitylating enzymes (DUBs) or shuttling ubiquitin (Ub) receptors. The main driving force for the conformational changes of 26S proteasome is provided by the AAA-ATPase via ATP binding and hydrolysis^15^. Therefore, the RPs undergo significant compositional and conformational changes to perform their biological functions.

A total of 22,000 particles for 26S proteasome were picked using a template matching method in pyTOM^43^ with a published *S. pombe* 26S proteasome map^47^ (EMDB accession code: EMD-2035) as the template. The picked subtomograms were extracted from the tomograms using RELION 3.0.8^4^. The rigid-body dynamics of the 26S proteasome were defined as the relative displacements between CP and two RPs, modelled as a three-body system. We performed structural heterogeneity analysis by training OPUS-DSD2 using subtomograms and their orientations directly from template matching results. The composition latent space was set to 12 dimensions and the dynamics latent space was set to 4 dimensions.

The composition latent space learned by OPUS-DSD2 was clustered into 20 classes. Classes 9 and 14∼19 contained clear densities for the 26S proteasome (**Fig.6a∼b**), and were selected for further clustering using KMeans (**Fig.6c∼d**). Focusing on one of the RP in 26S proteasome, pronounced compositional changes was observed in Rpn1—a ubiquitin receptor^48^, which were consistent with prior cryo-EM studies of the *S. pombe* 26S proteasome that reported occupancy variations in this subunit^47^. Specifically, the Rpn1 subunit is absent in class 12 but displayed progressively stronger and wider densities in classes 4, 15, and 9 (**Fig.6e**). Traversing principal component 5 (PC5) of the compositional latent space recapitulated these changes: the reconstruction at -1.1PC5 from OPUS-DSD2 revealed absence of Rpn1/2 and Rpt1/2, while the reconstruction at 0.2PC5 showed their emergence, and the reconstruction at 1.2PC5 exhibited enhanced density connectivity between Rpt1/2 and Rpn2 (**Fig.6f**).

**Figure 6.**
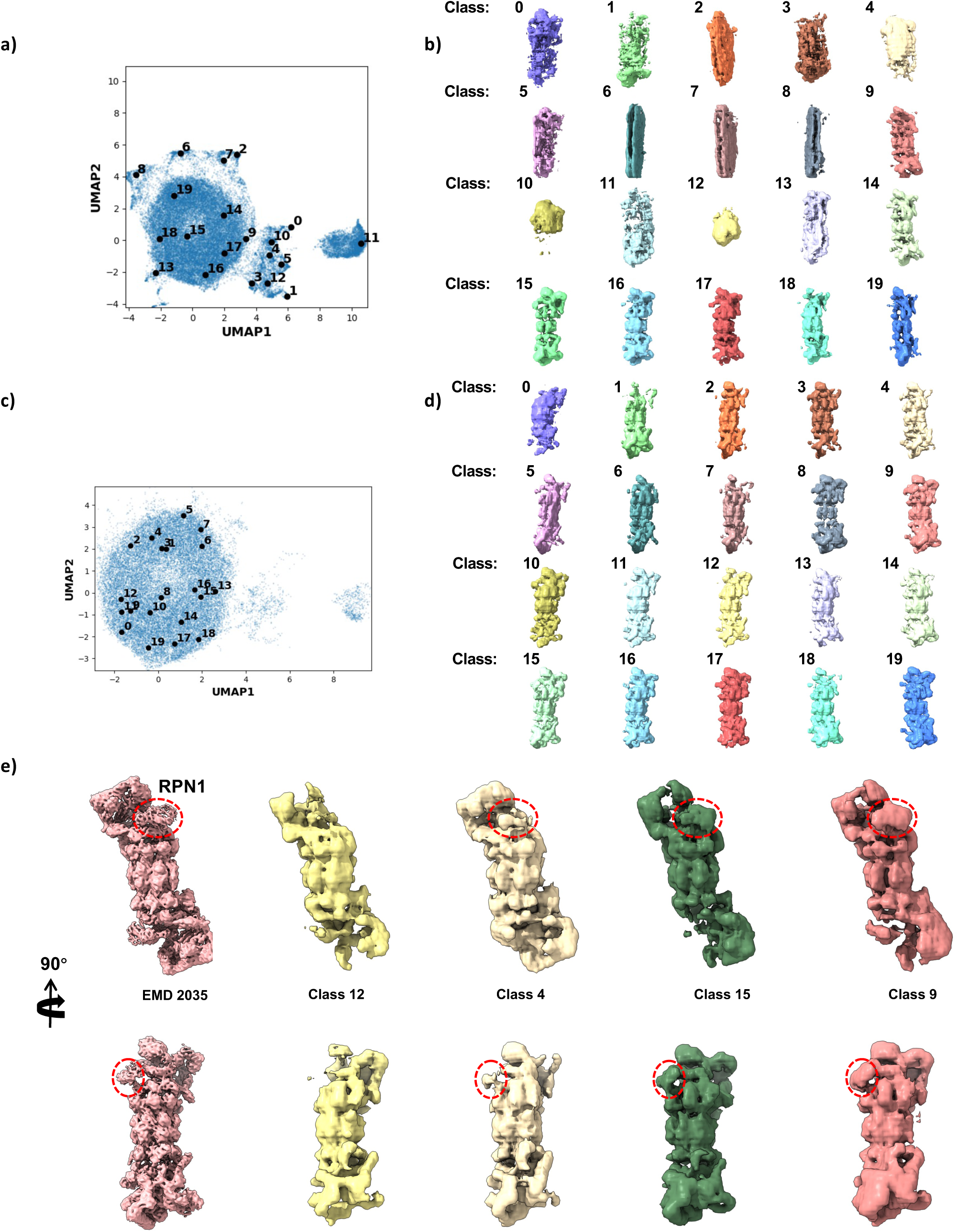
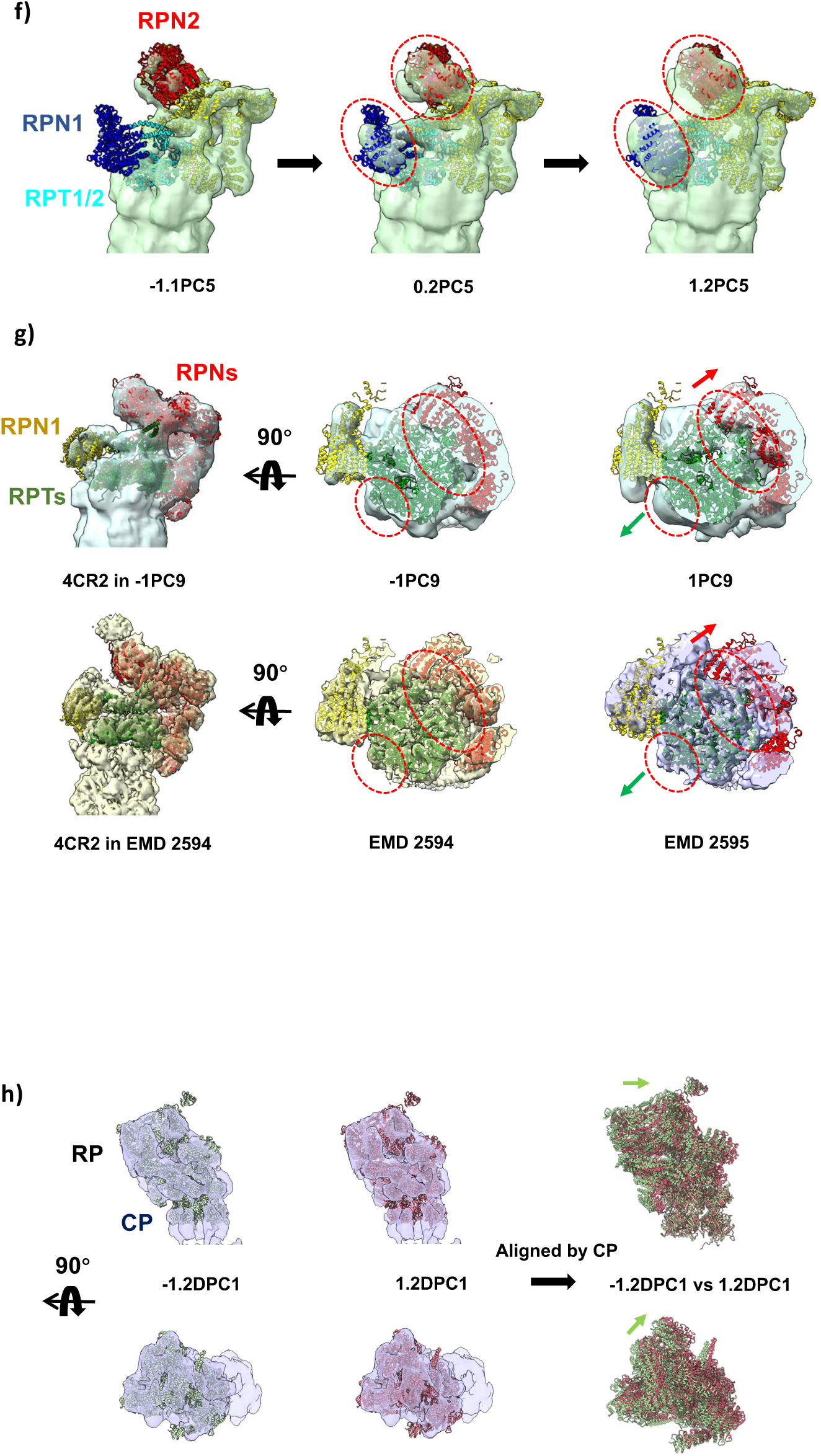
Heterogeneity analysis on *S. pombe* 26S proteasome (EMPIAR-10988). **a)**. UMAP visualization of the 12-dimensional composition latent space learned by OPUS-DSD2. Solid black dot represents the cluster center for labelled class. **b)**. Structures of 26S proteasome generated by the decoder of OPUS-DSD2 at the centroids of KMeans clusters in the composition latent space. **c)**. UMAP visualization of the 12-dimensional composition latent space learned by OPUS-DSD2 for classes 14∼19 in **a)**. Solid black dot represents the cluster center for labelled class. **d)**. Structures of 26S proteasome generated by the decoder of OPUS-DSD2 at the centroids of KMeans clusters in the composition latent space of classes 14∼19 in **a**). **e)**. Comparison between classes with different occupancies of RPN1 subunit. The sites showing densities for RPN1 subunits are marked by red dashed circles. A density map for *S. pombe* 26S proteasome (EMDB accession code: EMD 2035) is showing for reference. **f)**. Occuapacy changes of RPN1/2 and RPT1/2 subunits in density maps reconstructed by the Decoder of OPUS-DSD2 using latent codes at different locations along PC5 in composition latent space. The locations of latent codes are specified as xPCn. The sites showing densities for Rpn1/2 subunits are marked by red dashed circles. The atomic structure for *S. c* 26S proteasome (PDB code: 4CR2) was fitted into the density map reconstructed at 1.2PC5 for reference. The Rpn1 is colored in blue, the Rpn2 is colored in red, the Rpt1/2 are colored in cyan, the remaining Rpts are colored in yellow. **g)**. Conformational changes between AAA-ATPase and Rpns in density mpas reconstructed by the Decoder of OPUS-DSD2 using latent codes at different locations along PC9 in composition latent space. The atomic structure for *S. c* 26S proteasome (PDB code: 4CR2) was fitted into the density map reconstructed at -1PC9 and the density map EMD 2594 for reference. The Rpn1 subunit is colored in yellow, the ring-shaped AAA-ATPase formed by Rpts are colored in green, and the remaining Rpns are colored in red. Red dashed ellipses highlight the region of densities for Rpns and Rpts showing large displacements in relative to the atomic structures for reference states. The red arrows denote the direction of displacement for Rpns. The green arrows denote the direction of displacement for Rpts. **h**) Relative displacement between RP and CP in density maps reconstructed by the Decoder of OPUS-DSD2 using latent codes at different locations along DPC1 in dynamics latent space. The structure fitted with the density maps reconstructed by the Decoder of OPUS-DSD2 at the -1.2DPC1 is colored in green, and the structure fitted with the density maps reconstructed by the Decoder of OPUS-DSD2 at 1.2DPC1 is colored in red. The green and red structures in superposition are aligned w.r.t the CP. The green arrows denote the direction of displacement from the green structure to red structure.

Subtle conformational changes between the AAA-ATPase (Rpts) and Rpns can be resolved by traversing PC9 in the composition latent space. The atomic model for 26S proteasome (PDB code: 4CR2)^15^ rigidly fitted into the density map at -PC9 revealed outward displacement of Rpns and radial expansion of the Rpt ring when transiting to the state at PC9 (**Fig.6g**). Specifically, in 1PC9, the densities for Rpns are displaced outward compared to their original positions in -1PC9 indicated by the atomic model (**Fig.6g**). Moreover, the ring shape densities for Rpts are expanding compared to the original conformation represented by the green atomic model (**Fig.6g**). These movements mirror the s1-to-s2 conformational transition reported by Unverdorben *et. al.*^15^, as evidenced by comparable directional shifts in Rpns/Rpts between s1 (EMD accession code: 2594) and s2 (EMD accession code: 2595) (Fig.6g, **lower panel**). The atomic structure is fitted with the density map of s2 (Fig.6g, **lower panel**).

To analyze large-scale rigid-body dynamics between RPs and CP, DPC1 in the dynamics latent space was traversed using class 8 from the composition latent space as the template conformation. The atomic structure of the 26S proteasome (PDB code: 4CR2)^15^ can be well fitted into density maps reconstructed at -1.2DPC1 and 1.2DPC1 by rigid-body fitting, where the atomic structures of the RP and CP were fitted separately (Fig.6h **left panel**). Alignment of fitted atomic structures by CP revealed that RP translocates toward the center axis of CP when transiting from -1.2DPC1 to 1.2DPC1 (Fig.6h **right panel**), highlighting a coordinated motion potentially linked to substrate processing.

### *S. Pombe* Fatty Acid Synthase

Finally, we evaluated the performance of OPUS-DSD2 in the structural heterogeneity anlaysis of cryo-ET data for a sparsely populated specie, *Schizosaccharomyces pombe* fatty acid synthase (*S. Pombe* FAS) *in situ. S. Pombe* FAS, a type I fungal FAS, features two separate domes, each consisting of three β-subunits, and an equatorial wheel formed by six α-subunits^49^. The biosynthesis of fatty acids is performed by FAS inside the reaction chambers under domes, resulting in significant conformational flexibility in the barrel wall^49^.

A total of 10,000 particles for FAS were picked using template matching method in pyTOM^43^ with a published *S. c* FAS map^49^ (EMDB accession code: EMD-1623) as the template. The picked subtomograms were extracted from the tomograms using RELION 3.0.8^4^. The rigid-body dynamics of FAS was defined as the relative displacements among the upper dome, the equatorial wheel (formed by six α-subunits) and the lower dome. Given the *D*_3_ symmetry inherent to the FAS structure, we applied symmetry expansion to the pose parameters of subtomograms using RELION 3.0.8 as a preparatory step for structural heterogeneity analysis. Following this symmetry expansion, OPUS-DSD2 was trained on subtomograms with the assumption that their projection angles correspond to one of the six equivalent angles satisfying *D*_3_ symmetry.

The UMAP visualization of the 10-dimensional latent space learned by OPUS-TOMO shows distinct clusters, which we clustered into 20 classes by KMeans algorithm, and reconstructed 3D structure for each class by supplying cluster centroids into the trained Decoder of OPUS-DSD2 (**Fig.7a∼b**). Only class 14 shows complete densities for FAS, and contains 549 particles (**Fig.7b**). The class 14 was selected for further analyses by KMeans clustering and PCA (**Fig.7c∼d**).

**Figure 7.**
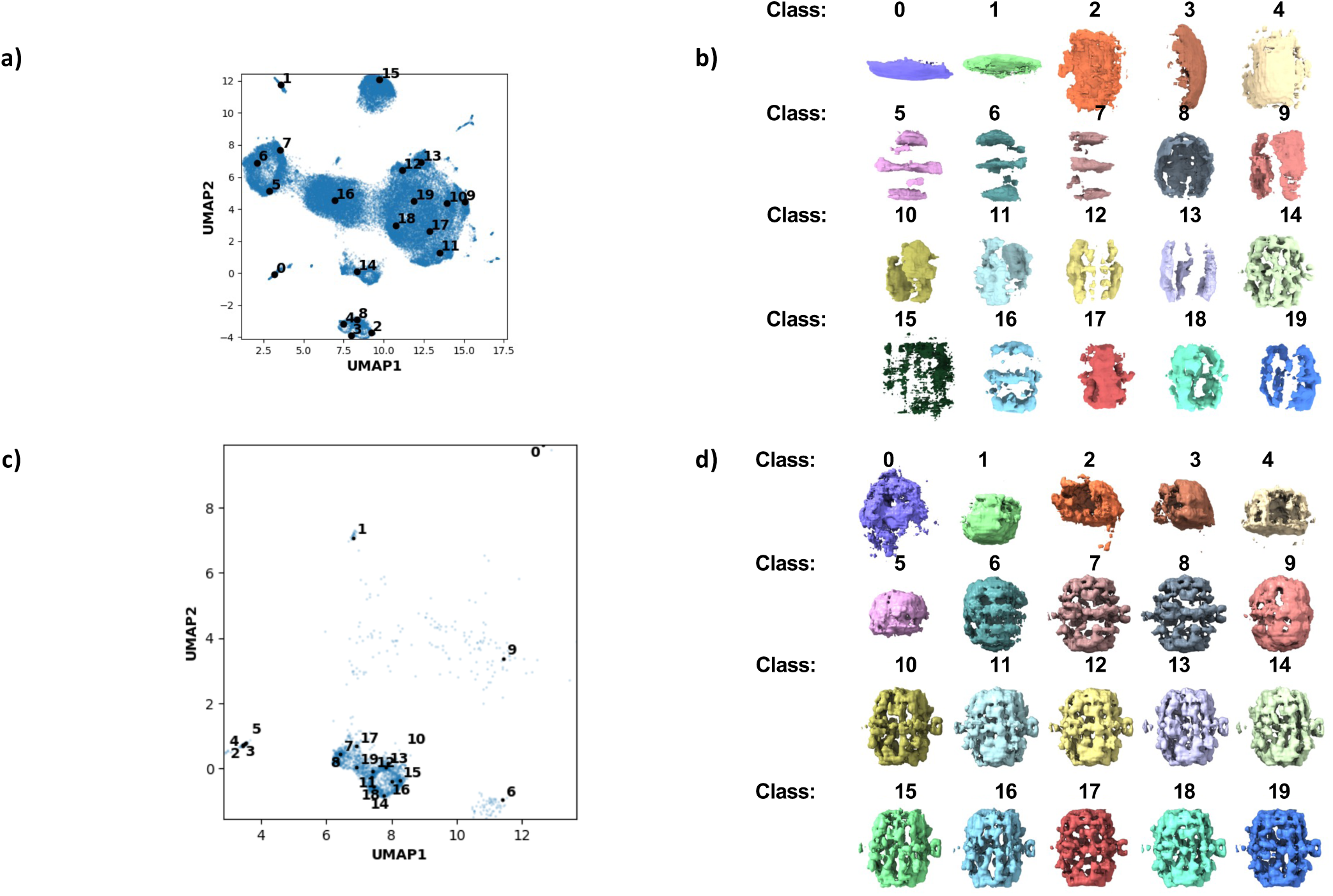
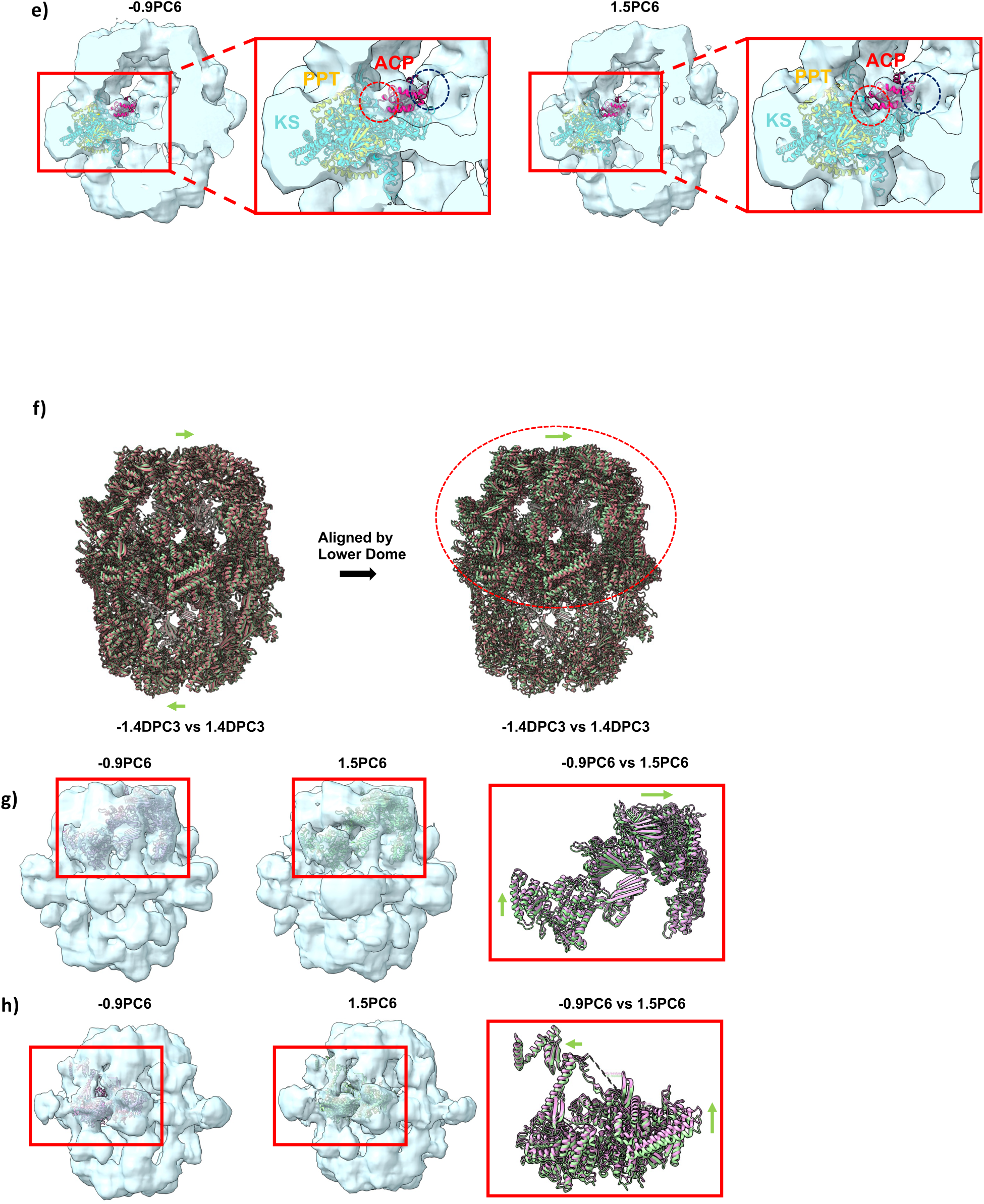
Heterogeneity analysis on *S. pombe* FAS (EMPIAR-10988). **a)**. UMAP visualization of the 10-dimensional composition latent space learned by OPUS-DSD2. Solid black dot represents the cluster center for labelled class. **b)**. Structures of *S. pombe* FAS generated by the decoder of OPUS-DSD2 at the centroids of KMeans clusters in the composition latent space. **c)**. UMAP visualization of the 12-dimensional composition latent space learned by OPUS-DSD2 for class 14 in **a)**. Solid black dot represents the cluster center for labelled class. **d)**. Structures of 26S proteasome generated by the decoder of OPUS-DSD2 at the centroids of KMeans clusters in the composition latent space of class 14 in **a**). **e)**. The movement of Acyl carrier protein (ACP) inside the reaction chambers revealed by density maps reconstructed by the Decoder of OPUS-DSD2 using latent codes at different locations along PC6 in composition latent space. The locations of latent codes are specified as xPCn. The atomic structure of *S. c* FAS (PDB code: 2VKZ) is fitted into PC6 for reference. The ACP is colored in red, the ketoacyl synthase (KS) dimer are colored in cyan, the phosphopantetheinyl transferase (PPT) domain is colored in yellow. Red dashed circles highlight the interface between ACP and KS dimer, yellow dashed circles highlight the interface between ACP and central hub. **f)**. The relative displacements of α subunit revealed by density maps reconstructed by the Decoder of OPUS-DSD2 using latent codes at different locations along PC6 in composition latent space. The atomic structure of an α subunit (PDB code: 2VKZ) were fitted into -0.9PC6 and 1.5PC6, respectively. The α subunit fitted with -0.9PC6 is colored in red, and the α subunit fitted with 1.5PC6 is colored in green. The green arrow indicate the direction of displacement from α subunit in 1.5PC6 to α subunit -0.9PC6. **g)**. The relative displacements of β subunit revealed by density maps reconstructed by the Decoder of OPUS-DSD2 using latent codes at different locations along PC6 in composition latent space. The atomic structure of an β subunit (PDB code: 2VKZ) were fitted into -0.9PC6 and 1.5PC6, respectively. The β subunit fitted with -0.9PC6 is colored in red, and the β subunit fitted with 1.5PC6 is colored in green. The red arrow indicates the direction of displacement from β subunit in 1.5PC6 to α subunit -0.9PC6. **h)**. The relative displacements of upper and lower domes revealed by density maps reconstructed by the Decoder of OPUS-DSD2 using latent codes at different locations along DPC3 in dynamics latent space. The atomic structure of *S. c* FAS (PDB code: 2VKZ) were fitted into -DPC3 and DPC3. The atomic structure fitted with -1.4DPC3 is colored in red, and the atomic structure fitted with 1.4DPC3 is colored in green. The atomic structures in the right panel were aligned w.r.t the lower dome. The green arrow indicates the direction of displacement from the upper dome of the atomic structure in 1.4DPC3 to -1.4DPC3. The red dashed ellipse marks the upper dome.

Acyl carrier protein (ACP) is a mobile domain inside the reaction chambers of type I fungal FAS^49^. Traversing PC6 of the composition latent space learned by OPUS-DSD2 reveals the movement of ACP docked to the ketoacyl synthase (KS) dimer (**Fig.7e**). By fitting the atomic model of type I FAS (PDB code: 2VKZ)^50^ into the density map reconstruct at 1.5PC6, the we observed ACP domain is disengaging from the barrel wall part of KS at -0.9PC6 and relocates towards the central hub (**Fig.7e**).

The latent spaces learned by OPUS-DSD2 also encode dynamics of FAS at different scales. Firstly, the dynamics latent space is traversed to reveal rigid-body dynamics between domes. The atomic structure for FAS (PDB code: 2VKZ)^50^ was fitted into the density maps reconstructed by OPUS-DSD2 using latent codes at -1.4DPC3 and 1.4DPC3 in dynamics latent space. Rigid-body fits of different domes were separately performed. Superposition of fitted atomic models revealed displacements of two domes in opposite directions (Fig.7f **left panel**). After aligning fitted atomic models by the lower dome, a pronounced twist of the upper dome relative to the lower dome was evident when transitioning from -1.4DPC3 to 1.4DPC3 (Fig.7f **right panel**). Secondly, subunit-specific rearrangements can be revealed by traversing the composition latent space. The atomic structures of α and β subunits were separately fitted into the density maps reconstructed by OPUS-DSD2 using latent codes at -PC6 and PC6 in composition latent space. By the superposition of the β subunits fitted with conformations at - PC6 and PC6, the top part of β subunit is contracting towards the center of reaction chamber when transiting from -0.9PC6 to 1.5PC6 (**Fig.7g**). Meanwhile, the superposition of the α subunits fitted with conformations at -0.9PC6 and 1.5PC6 shows that α subunit is shifting downwards when transiting from -0.9PC6 to 1.5PC6 (**Fig.7h**).

## Discussion

In this paper, we present OPUS-DSD2, a unified framework for resolving structural heterogeneities in both cryo-EM and cryo-ET data. OPUS-DSD2 incorporates an MLP-based rigid-body dynamics model to explicitly capture dynamics inherent in experimental datasets. Through extensive validations on cryo-EM datasets and *in vivo* cryo-ET samples, we demonstrate that OPUS-DSD2 significantly enhances the reconstruction quality of its decoders, improves the interpretability of structural heterogeneity analyses, and resolves biologically relevant heterogeneities – even when applied to cryo-ET tomograms using only template matching alone.

OPUS-DSD2 has shown the ability to disentangle structural heterogeneity into orthogonal modes of variations. Experiments on real cryo-EM datasets using OPUS-DSD2 demonstrate that principal components of the dynamics latent space model inter-subunit rigid-body movements, while those in composition latent space capture localized structural rearrangements. This separation facilitates intuitive interpretation of complex structural changes. OPUS-DSD2 achieves this interpretability through the new deformable volume representation, which combines a 3D structural map with a subunit-specific deformation field. The 3D volume provides a holistic representation of structural variability, while the deformation field facilitates the modelling of rigid-body dynamics of individual subunits using much less parameters. Unlike the multi-body dynamics model proposed in RELION^28^, OPUS-DSD2 employs a continuous blending of subunit movements near boundaries, ensuring physically plausible, smooth transitions between adjacent subunits. This hybrid representation allows the 3D convolutional network to focus on intra-subunit structural changes. OPUS-DSD2 effectively decomposes the complex’s overall dynamics^27–29^ and structural changes in different scales, and associates them with different principal components of latent spaces, greatly increasing the interpretability of the structural heterogeneity resolving results.

The improved reconstruction quality of OPUS-DSD2’s decoder stems from synergistic interactions between its dynamics decoder and composition decoder. By explicitly modeling subunit displacements via the deformation field, the composition decoder can represent diverse conformations using a single latent code, effectively pooling information across particles with varying dynamics. This equips the 3D convolutional network with equivariance to rigid-body motions, enabling supervision from a broader ensemble of particles and enhancing reconstruction accuracy. Conversely, higher-quality 3D volumes from the composition decoder improve the precision of subunit displacement estimates by the dynamics decoder, creating a positive feedback loop.

A challenging task is resolving structural heterogeneity for cryo-ET data. Cryo-ET captures *in situ* snapshots of diverse macromolecular species, offering unprecedented potential to elucidate biological mechanisms by linking molecular-scale structural changes to function. Traditional workflows of cryo-ET, however, involves laborious multi-step processes: template matching to identify target complexes in crowded cellular environments, followed by iterative 3D classification and subtomogram averaging to resolve functional states^51^. These workflows are computationally intensive and impractical for proteome-scale studies. A transformative advance would be single-shot reconstruction of distinct functional states directly from template-matching results—a task hindered by three key challenges. The first challenge is pose assignment errors. The orientation determined by template matching often suffers from inaccuracies due to coarse angular sampling and conformational mismatches between particles and consensus reference models. Meanwhile, template matching totally omits translational alignment. The errors in misassigned poses propagate into downstream reconstructions, distorting inferred conformations. Accurate reconstruction requires precise alignment relative to native structures. The second challenge is non-specific particle selection. Low-resolution templates may inadvertently pick various macromolecular species in noisy tomograms. The third challenge is intrinsic structural heterogeneity. Target complexes exhibit compositional and dynamical variations reflecting their functional states *in situ*.

OPUS-DSD2 addresses these challenges, enabling robust heterogeneity analysis using raw template-matching outputs. Its dynamics model treats pose errors as a manifestation of rigid-body dynamics where the whole macromolecule acts as a rigid body, i.e., the movements of subunits become highly degenerate. The pose errors are encoded into the dynamics latent space. For example, traversals along principal components of this space for S. pombe 80S ribosomes, 26S proteasomes, and fatty acid synthase (FAS) reveal that OPUS-DSD2 successfully captures global rotational/translational displacements of entire complexes. Concurrently, distinct macromolecular species picked during template matching are disentangled as separable clusters in the composition latent space. This dual latent representation allows OPUS-DSD2 to reconstruct high-fidelity density maps of target complexes and resolve transient functional states—even from subtomograms with erroneous initial poses—providing mechanistic insights into their biological roles in cellular contexts.

In conclusion, the introduction of OPUS-DSD2 offers a neural-network-based paradigm in cryo-EM/ET data analysis, and allows for explicit modelling of dynamics and improved structural heterogeneity analysis. Additionally, OPUS-DSD2 can leverage the results of structural heterogeneity analysis produced by OPUS-DSD, which can uncover crude patterns of macromolecular movements in cryo-EM data. OPUS-DSD2 can employ these preliminary modes to define the partition of rigid subunits and their rotation axes in dynamics, iterating to refine these results further. However, as with any novel method, it is necessary to verify its performance with a broad range of macromolecular complexes and under different experimental conditions. This, in turn, will provide a rigorous assessment of its generalizability and robustness.

## Methods

### Notations

*x*, ***x*** and *X* will be denoted as scalars, vectors and matrices, respectively. ‖·‖ represents the Euclidean norm of a tensor. χ denotes the Fourier transform of the multidimensional array *X*.

### Deformation field

OPUS-DSD2 represents a 3D conformation by explicitly modelling its composition and deformation. The composition of a 3D conformation can be represented by a volume on a discrete 3D grid, which is referred as template, and the deformation in relative to the template can be represented by a 3D deformation field. Each voxel in the deformation field records its original position in the template volume. The 3D conformation in OPUS-DSD2 is sampled from the template according to the deformation field. Formally, the 3D conformation can be given by the composition of two functions *V*(***x*** + ***g***(***x***)): ℝ^3^ → ℝ, where *V*(·) is the template which maps a grid point ***x*** to a voxel value, and ***g***(***x***): ℝ^3^ → ℝ^3^ is the deformation field which record the displacement between the voxel ***x*** in the deformed conformation and its source voxel ***x*** + ***g***(***x***) in template. Next, we give a detailed definition about the deformation field.

In OPUS-DSD2, the deformation of macromolecule is assumed to be compromised of individual movements of multiple subunits. The 3D deformation field is constructed through several steps. Firstly, a 3D consensus model is segmented into a set of pre-defined subunits, each of which has a specific shape with masses inside and undergoes rigid-body movement. Mathematically, the subunit can be defined as a function on the 3D grid, which is of the form

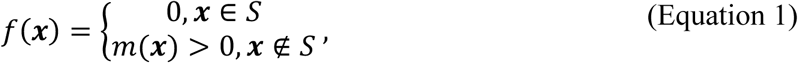

where *S* refers to the set of grid points belongs to the subunit, and *m*(***x***) is the mass at point ***x***. We can further define some statistics about the subunit. For a subunit with center ***c***_*i*_ ∈ ℝ^3^, and suppose its principal axes form a matrix *P*^*i*^, where each row is one principal axis, its semi-principal diameters can be defined as

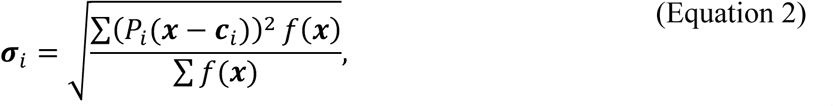

where the summations are over all grid points in the 3D grid. For a grid point ***x*** = {*x*, *y*, *z*} ∈ ℝ^3^ in a subunit *i*, suppose the subunit undergoing a rigid-body displacement where the subunit is rotated by a rotation matrix *R_i_* ∈ *SO*(3), and translated by the translation vector ***t**_i_* ∈ ℝ^3^, let the center of mass (COM) of the subunit be ***c**_i_*, the rigid-body displacement of this point can be expressed as

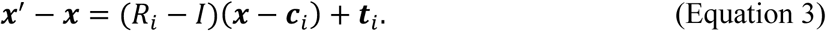

For a grid point in a large complex which is comprised of multiple subunits, each of which undergoes independent movement, its displacement should be the combination of those movements. We propose to blend the movement from a subunit at a grid point according to its distance w.r.t the center of subunit. Specifically, for a grid point ***x***, suppose the principal axes of subunit *i* form a matrix *P_i_* as preceding, the displacement incurred by the movement of subunit *i* decays as,

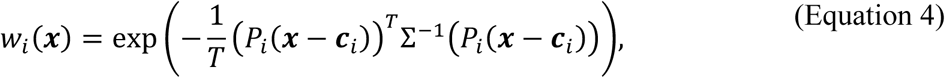

where *xj* is the *j*th component of the grid point ***x***, Σ is a diagonal matrix with its diagonal element Σ*_jj_* = σ*_i,j_*, i.e., the semi-principal diameter of subunit *i* along the *j*th axis, and *T* is a constant for controlling the falloff of the weight function. The weight function should be normalized if there are *n* subunits in the complex. The normalized weight function can be expressed as,

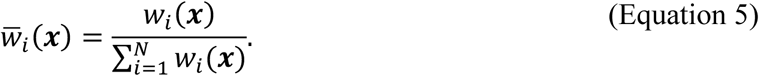

In a complex with *n* subunits, the displacement of a grid point is the linear combination of movements of different subunits, which can be written as,

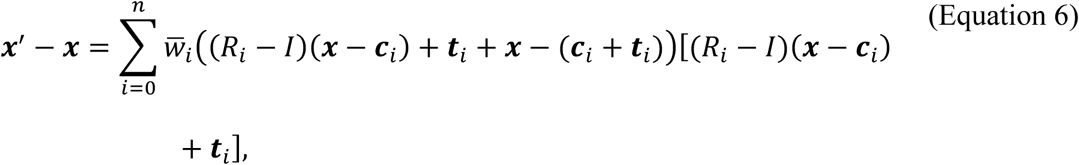

where 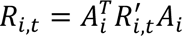 controls the scale of the movement of subunit *i*. We hence defined the deformation field ***g***, with which the deformed grid can be represented as ***x***^+^ = ***x*** + ***g***(***x***). In OPUS-DSD2, the rigid-body transformation parameters, *R_i_* and ***t**_i_*, are predicted by the neural network. They should be restrained to rule out unrealistic deformations. We assume that the translational modes of subunits are generated by rotating COMs of subunits in relative to the COM of a common fixed subunit, i.e., ***t****_i_* = (*R_i,t_* – *I*)(***c**_i_* – ***c***_f_), where the COM ***c**_i_* is rotated by *R*_),4_ in relative to COM ***c**_j_*, and the translation of subunit can be obtained as the difference between the rotated COM and the original COM. The translation defined in this way retains the distance between COMs of subunits. The determination of the translation of each subunit comes down to estimating two rotation matrices. We can further limit the range of motion by restraining the rotation matrix to be an identity matrix, which inevitably hurts the expression power of the deformation model. To maximize the capacity of the deformation model under the identity restraint, the rigid-body movement should be defined in an appropriate reference frame, i.e., the *z* axis should align with the rotation axis. The rotation matrix which generates the translation of COM of subunit can be decomposed as 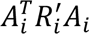, where the rotation matrix *A_i_* aligns the rotation axis of the COM of subunit to the *z* axis, and 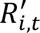 is the rotation matrix predicted by neural network. The rotation matrix of the subunit can be decomposed similarly as *R_i_* = 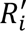, where 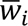 is the rotation matrix predicted by neural network. Our method adopts quaternion to represent rotation. The neural network in decoder outputs the last three dimensions of the quaternion, while the first dimension of quaternion is fixed to be a positive value, 8, which can regularize it to be an identity rotation.

### Image Formation Model

Cryo-EM and cryo-ET are two related techniques with very similar image formation models. We start by defining the image formation model for cryo-EM with which the image formation model for cryo-ET can be easily defined. The images collected by cryo-EM are 2D projections of a 3D molecular structure which undergoes a series of transformations from a template 3D volume. For a molecular structure *V*′, let the template volume be *V*, suppose *V*′ is rotated from *V* by rotation matrix *R_t_* and deformed by the deformation field ***g***, the voxel in the 3D molecular structure has the following relation w.r.t the voxel in the template volume,

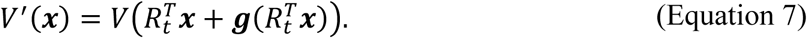

The 3D molecular structure can then be sampled from the template volume using spatial transformer. The 2D projection of the molecular structure can be obtained by summation along the *Z* axis, namely,

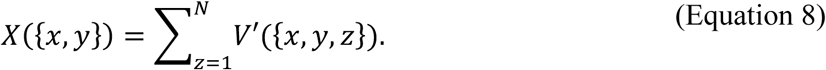

In real cryo-EM experiments, the images are not in perfect focus, the objective lens presents aberration and there are significant amount of noises when recording images. Hence, the 2D projections are should be corrupted by the contrast transfer function which describes the effects of underfocus and the aberration of objective lens. In Fourier domain, let the Fourier transform of 2D projection be χ, the Fourier transform of the experimental image χ′ is of the form^52^

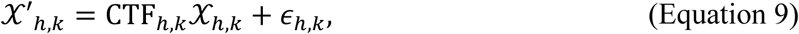

where CTF_*h,k*_ is the contrast transfer function at frequency vector [*h, k*], and ∊_*h,k*_ is the noise introduced during experiment. To reconstruct a 3D tomogram for the cellular sample, cryo-ET further collects a series of 2D projections with different tilt angles. Therefore, the subtomogram in cryo-ET is of three dimensionality and consists of 2D images with different tilt angles. Similarly, in Fourier domain, let the Fourier transform of 3D structure be *V*, the Fourier transform of the subtomogram χ′ can be written as,

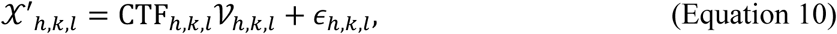

where CTF*_h,k,l_* is the contrast transfer function at frequency vector [*h, k, l*], and ∊*_h,k,l_* is the noise. The CTF model for the subtomogram in cryo-ET hence becomes 3D and accounts for the CTF of each tilt image. It can be obtained by backprojecting the 2D CTFs of all images in the tilt series into an empty 3D volume in Fourier domain. Each 2D CTF is placed as a central slice in the 3D CTF, whose orientation is determined by its tilt angle. Reconstructing the 3D CTF model for subtomograms by backprojection naturally handles the missing wedges as the region which is not covered by tilt series remains empty. Consequently, with the introduction of 3D CTF, the image formation model for cryo-EM in (Equation 9) can be viewed as a special case of the image formation model given in (Equation 10), where the 3D CTF has only one nonzero 2D slice. Henceforth, we will use the 3D image formation in (Equation 10) for defining training objective and refer to all experimental image from cryo-EM/ET as subtomogram.

We hence define an image formation model to generate subtomogram from a template volume according the specified deformation. It is fully differentiable and enables us to learn the template volume and deformation field simultaneously.

### Training Objective

The main objective of OPUS-DSD2 is learning the template volumes and deformation fields using subtomograms. In OPUS-DSD2, two latent spaces are designated to encode the template volume and deformation field, respectively. Two neural networks generate these objects conditioned on the corresponding latent codes. Let the neural network that represents the template volume be *V*, and the neural network that represents the deformation field be ***g***, for a particle, the subtomogram *X* reconstructed by these two neural networks is of the form,

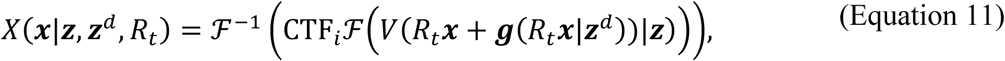

where ***x*** represents the coordinate of a voxel, ***g*** is the composition latent code of the particle for the network *V*, ***g**^d^* is the dynamics latent code for the network ***g***, CTF_i_ represents the 3D CTF for the particle, and ℱ and ℱ^-.^ denote Fourier transform and inverse Fourier transform, respectively. The latent spaces and neural networks can be trained by minimizing the difference between the image reconstructed by the image formation model and the experimental subtomogram, that is,

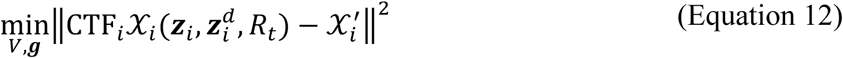

where ***g**_i_* is the latent code for the template volume of image *i*, and 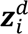 is the latent code for the deformation field of image *i*. However, the latent codes ***g***_)_ and 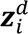 are unknown. In practice, to efficiently train the content generation neural networks and guarantee the smoothness of latent space, variational autoencoder (VAE) architecture is adopted. The distributions of latent codes ***g*** and ***g**^d^* are inferred by an encoder network *f*, and their distributions are restrained to be close to the standard gaussian distribution. In OPUS-DSD, we proposed a structural disentanglement prior to enforce the smoothness of latent space of template volumes^18^. Similarly, OPUS-DSD2 leverages a combination of the training objective of β-VAE^36^ and the structural disentanglement prior to learn a smooth latent space for representing template volume. More specifically, let the dimension of 3D volume be *N*^3^, the training objective of OPUS-DSD2 is of the form,

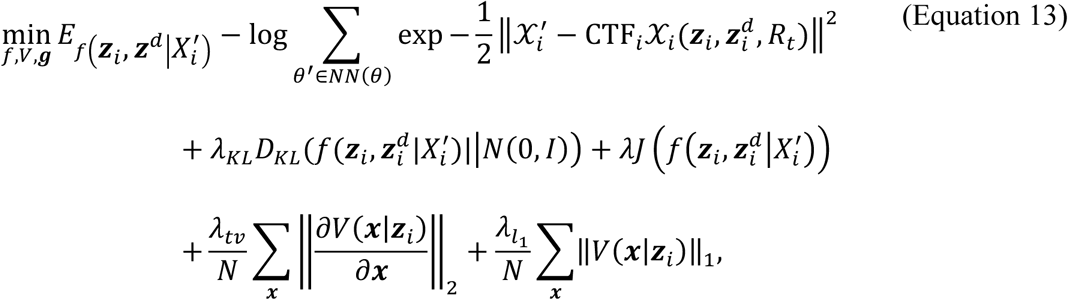

where the first term is expectation of error between the experimental subtomogram 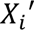 and the reconstruction over the distribution of latent codes, *N*(0, *I*) is the standard gaussian distribution, *D_KL_* is the KL divergence between the distributions of latent codes given by the encoder network *f* and the standard gaussian distributions, λ*_KL_* is the restraint strength for the KL divergence, λ is the restraint strength of structural disentanglement prior for the latent codes, λ*_l_*_1_ is the restraint strength for the total variation of the template volume, and λ*_tv_*, represents the restraint strength of the *L*_1_ norm of the template volume. It is also worth noting that the expectation term in (Equation 13) is intractable, and is approximated by the reparameterization trick during training ^35^.

### Training

Our networks are trained end-to-end using only subtomograms. The starting point of training is a set of 2D images after consensus 3D refinement in RELION^7^ or cryoSPARC^34^, or 3D subtomograms picked by template matching in pyTOM^8^. The pose parameters of inputs are determined w.r.t to a consensus reference model. Before training, a specific fraction of subtomograms is randomly assigned to the training set which is used to train the neural networks, while the remaining subtomograms are reserved for the validation set which solely serves to evaluate the reconstruction quality of neural network. A customized batching process is designed to approximately compute the structural disentanglement prior. The subtomograms are classified into 48 different projection classes according to the first two Euler angles of their rotation w.r.t the consensus model using HEALPix^53^. During training, a projection class is randomly chosen, and a batch of subtomograms within the projection class is sampled for training. This batching process guarantees the subtomograms in a batch come from the same projection class, and approximates the intra-class comparison term of latent codes in the structural disentanglement prior. A sampled batch of subtomograms undergo a series of data augmentation operations which was defined in OPUS-DSD^18^. The augmented subtomograms are fed into the encoder network to compute the means and standard deviations of their composition and dynamics latent codes. We sample latent codes according to computed distributions and supply them into the composition and dynamics decoders, which generate template volume and deformation field, respectively. The final volume for the input image is reconstructed by deforming the template volume according to the deformation field. The deformed volumes are projected into 2D reconstructions following the image formation model which is described before. The reconstructions together with input images without augmentation form the reconstruction loss function. We also added the priors for latent codes, and the total variation^54^ and *L*_1_ norms^55^ for the template volumes with specified weights to the training objective according to (Equation 13). The total loss function is optimized by the Adam optimizer^56^.

### Hyperparameters’ settings

It is of critical importance to setting suitable hyperparameters for training OPUS-DSD2 since they balance input fitting and latent space regularization. Suitable hyperparameters can guarantee the quality of heterogeneity discovery. Since reconstruction losses are computed over a batch of subtomograms with high level of noises, while the priors are computed over the denoised signals, the reconstruction losses are much noisier than the priors. To balance the variances of reconstruction losses and the priors while avoiding posterior collapse^57^, the scale of hyperparameters should be proportional to the SNRs of cryo-EM datasets. Moreover, the SNRs of cryo-EM/ET datasets are varying due to the masses of macromolecules and experimental setup. The hyperparameters can be conveniently set on an absolute scale considering the SNR of dataset. Using a batch of *N* subtomograms, the SNR can be estimated as,

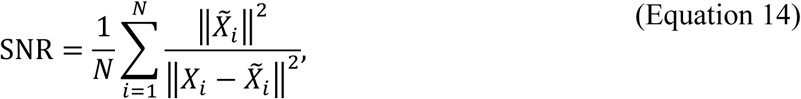

where 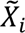 is the subtomogram reconstructed by OPUS-DSD2, and *X_i_* is the experimental subtomogram. 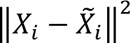 estimates the norm of noises in the experimental subtomogram, while 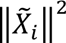 estimates the norm of signals. The restraint strength for KL divergence λ is then set in relative to the estimated SNR, and of the form λ × SNR.

We fixed the restraint strength for the total variation of 3D volume λ*_tv_* = 0.5 and the restraint strength for LASSO λ*_l_*_1_ = 0.15. For cryo-EM data, the learning rate was set to 10^-H^ that was decayed by 0.95 at each epoch for all experiments. For cryo-ET data, the learning rate was set to 3 × 10^-I^ that was decayed by 0.95 at each epoch for all experiments. The batch size was set to 64 during the training. All trainings were performed on 4 Nvidia V100 GPUs. The number of samples for interclass comparison was set to 12800. The numbers of kNN for interclass or intraclass comparison were set to 128 or 4, respectively, while the number of kFP for interclass comparison was set to 512. The latent codes of particles were updated with a momentum of 0.7.

## Acknowledgements

The research was partially supported by National Key Research and Development Program of China (No.2021YFF1200400), Shanghai Municipal Science and Technology Major Project (No.2018SHZDZX01) and ZJLab. The authors specially thank James Krieger for his contribution to the code improvement of OPUS-DSD2.

## Author Contributions

Z.L., Q.W., and J.M. conceived the work. Z.L. designed the algorithm and implemented the software. Z.L. and X.C. performed the experiments. All authors wrote the final manuscript.

## Supplementary Video Legends

**Supplementary Video 1.** Traversal of the third and fourth principal components of the composition latent space of the pre-catalytic spliceosome corresponds distinct compositional changes between Helicase and Foot.

**Supplementary Video 2.** Traversal of the first principal component of the dynamics latent space of the pre-catalytic spliceosome reveals the concerted folding of SF3b and Helicase onto the Core.

**Supplementary Video 3.** Traversal of the second principal component of the dynamics latent space of the pre-catalytic spliceosome reveals the folding of Helicase onto the Core and horizontal swing of the SF3b.

**Supplementary Video 4.** Traversal of the first and the third principal components of the composition latent space of Pf80S ribosome reveal different compositional changes.

**Supplementary Video 5.** Traversal of the second principal component of the dynamics latent space of the Pf80S ribosome reveals the relative rotation between the SSU and LSU.

**Supplementary Video 6.** Traversal of the first and the second principal components of the dynamics latent space of S trimer reveal two orthogonal modes of movements.

**Supplementary Video 7.** Traversal of the third principal component of the composition latent space of S trimer reveals the open and close of the S1 NTD.

